# A cholinergic spinal pathway for the adaptive control of breathing

**DOI:** 10.1101/2025.01.20.633641

**Authors:** Minshan Lin, Giulia Benedetta Calabrese, Anthony V. Incognito, Matthew T. Moore, Aambar Agarwal, Richard J.A. Wilson, Laskaro Zagoraiou, Simon A. Sharples, Gareth B. Miles, Polyxeni Philippidou

**Author notes:** Correspondence to: Polyxeni Philippidou Gareth B. Miles Simon A. Sharples. co-first authors (Minshan Lin, Giulia Benedetta Calabrese). co-senior authors (Simon A. Sharples, Gareth B. Miles, Polyxeni Philippidou).

## Abstract

The ability to amplify motor neuron (MN) output is essential for generating high intensity motor actions. This is critical for breathing that must be rapidly adjusted to accommodate changing metabolic demands. While brainstem circuits generate the breathing rhythm, the pathways that directly augment respiratory MN output are not well understood. Here, we mapped first-order inputs to phrenic motor neurons (PMNs), a key respiratory MN population that initiates diaphragm contraction to drive breathing. We identified a predominant spinal input from a distinct subset of genetically-defined V0_C_ cholinergic interneurons. We found that these interneurons receive phasic excitation from brainstem respiratory centers, augment phrenic output through M2 muscarinic receptors, and are highly activated under a hypercapnia challenge. Specifically silencing cholinergic interneuron neurotransmission impairs the breathing response to hypercapnia. Collectively, our findings identify a novel spinal pathway that amplifies breathing, presenting a potential target for promoting recovery of breathing following spinal cord injury.

---

Motor circuits of the nervous system must operate over a dynamic range to adapt their output and generate behaviors of varying intensities in response to environmental challenges. This is especially critical for breathing, a motor behavior essential for maintaining blood gases required to sustain the metabolism of vital organs, including the brain, heart, and kidneys^1^. To meet metabolic demands during physiological challenges such as exercise or the changing environmental conditions associated with altitude, the neuronal network that controls breathing must operate over a large dynamic range. Specialized circuits have evolved to support robust, yet adaptable breathing in terrestrial vertebrates. While a network located in the brainstem is largely responsible for generating the rhythm and pattern of breathing^2–4^, motor neurons (MNs) projecting to muscles in the periphery are the final output of respiratory circuits. In mammals, phrenic motor neurons (PMNs) located in the cervical spinal cord innervate the diaphragm, the major inspiratory muscle that drives airflow into the lungs. The strength of diaphragmatic contraction determines the volume of each breath. Thus, precise regulation of PMN activity is vital for aligning breathing with metabolic needs and ensuring appropriate responses to environmental challenges.

Both central and peripheral chemoreflexes safeguard CO_2_/pH homeostasis by mediating rapid ventilatory responses to atmospheric gas fluctuations and changes in metabolic demands^5^. Inputs from diverse sources, including central and peripheral CO_2_ and O_2_ chemoreceptors, are integrated by brainstem networks to generate multidimensional changes in ventilation^6–13^. Elevated brain CO_2_ (hypercapnia), for example, leads to a significant increase in both the respiratory frequency and amplitude of diaphragm contractions. While increases in frequency are mediated by projections from chemoreceptor neurons to key rhythmogenic regions confined to the brainstem^9,14^, mechanisms for increasing breathing intensity likely involve direct modulation of PMN activity and increased drive to PMNs from distinct pre-motor populations.

While the brainstem circuits that underlie the generation and modulation of the breathing rhythm have been well defined^2–4,15–18^, much less is known about the topography, molecular identity, and function of downstream neurons that directly project to PMNs. Antidromic stimulation and retrograde tracing has revealed monosynaptic connections to PMNs, primarily from the excitatory rostral ventral respiratory group (rVRG) in the brainstem, which drive PMN activation during inspiration^19–25^. Although direct PMN inputs from spinal interneurons have been identified^20,23,26–28^, the degree of connectivity between PMNs and local interneurons is unclear. Recent rabies virus tracing experiments in neonatal mice suggested that the number of direct PMN inputs that originate in the spinal cord is minor^25^; however, tracing experiments with polysynaptic pseudorabies viruses (PRV) in adult rats, cats, and ferrets reveal more extensive spinal respiratory circuits^20,21,23^. Multielectrode array recordings from the cervical spinal cord also identified interneurons with respiratory-related activity that are synaptically coupled to PMNs^29,30^, and excitatory interneurons at cervical and thoracic levels are able to sustain breathing after spinal cord injury (SCI)^31–33^. Despite their roles in promoting respiratory recovery following SCI, we know little about the functions of spinal interneurons, including genetically-defined subsets, in the control of breathing^34–37^. One possibility is that local spinal circuits may be important for controlling breathing intensity through direct modulation of respiratory MN output. We therefore hypothesized that distinct classes of spinal interneurons may regulate PMN activity, providing a hierarchically arranged gain control system for breathing that is spatially segregated from the brainstem-derived rhythm generator.

Here, we combined a genetic strategy with rabies virus-mediated tracing to label neurons with monosynaptic inputs to PMNs. We identified a morphologically and topographically distinct population of Pitx2+ V0_C_ interneurons, largely localized in the cervical spinal cord, with substantial direct projections to PMNs. We find that these interneurons are functionally integrated into respiratory circuits, can modify PMN output through M2 muscarinic receptors, and are activated in response to CO_2_ exposure. Finally, we show that inhibiting cholinergic neurotransmission from Pitx2+ interneurons impairs the response to hypercapnia. We propose that spinal cholinergic interneurons provide a novel node for breathing gain control that is spatially segregated from the brainstem and can be recruited to amplify breathing under high metabolic demands and intense respiratory challenges. These interneurons represent an accessible therapeutic target for conditions in which respiratory function is compromised, such as SCI or Amyotrophic Lateral Sclerosis (ALS).

## Results

### PMNs receive substantial monosynaptic inputs from the spinal cord

To investigate the contribution of spinal networks to PMN activity, we combined a genetic strategy with rabies-based monosynaptic retrograde tracing to map inputs to PMNs. We utilized a modified glycoprotein (G protein)-deleted mCherry-tagged rabies virus (RabiesΔG-mCherry). Typically, the rabies virus requires G protein for transsynaptic transport, which is not endogenously expressed in mammalian cells. Therefore, G protein deletion from the rabies virus ensures that transsynaptic transport is not possible from the virus itself in wildtype mammalian neurons. To activate the transsynaptic transport mechanism and enable monosynaptic labeling specifically from MNs, we crossed *RphiGT* mice, which express G protein after Cre-mediated recombination, to *Choline acetyltransferase* (*ChAT)::Cre* mice (*ChAT::Cre; RphiGT*) to induce G protein expression only in cholinergic neurons, which include MNs (Extended Data Fig. 1A)^38^. We validated the expression of G protein in *ChAT::Cre; RphiGT* mice by *in situ* hybridization at postnatal day (P)4. Notably, despite the existence of cholinergic interneurons in the spinal cord, the expression of G protein was only detectable in MNs at P4 (Extended Data Fig. 1B). We injected RabiesΔG-mCherry unilaterally into the diaphragm, which is solely innervated by PMNs, of *ChAT::Cre; RphiGT* mice at P4 to label PMNs and trace their synaptic inputs (Fig. 1A). Our viral injections specifically targeted PMNs, as seen by the absence of labeled ventral roots and MNs at thoracic and lumbar levels of the spinal cord (Extended Data Fig. 1C). Each injection labeled 1-5 starter PMNs (Extended Data Fig. 2A and B).

**Figure 1.**
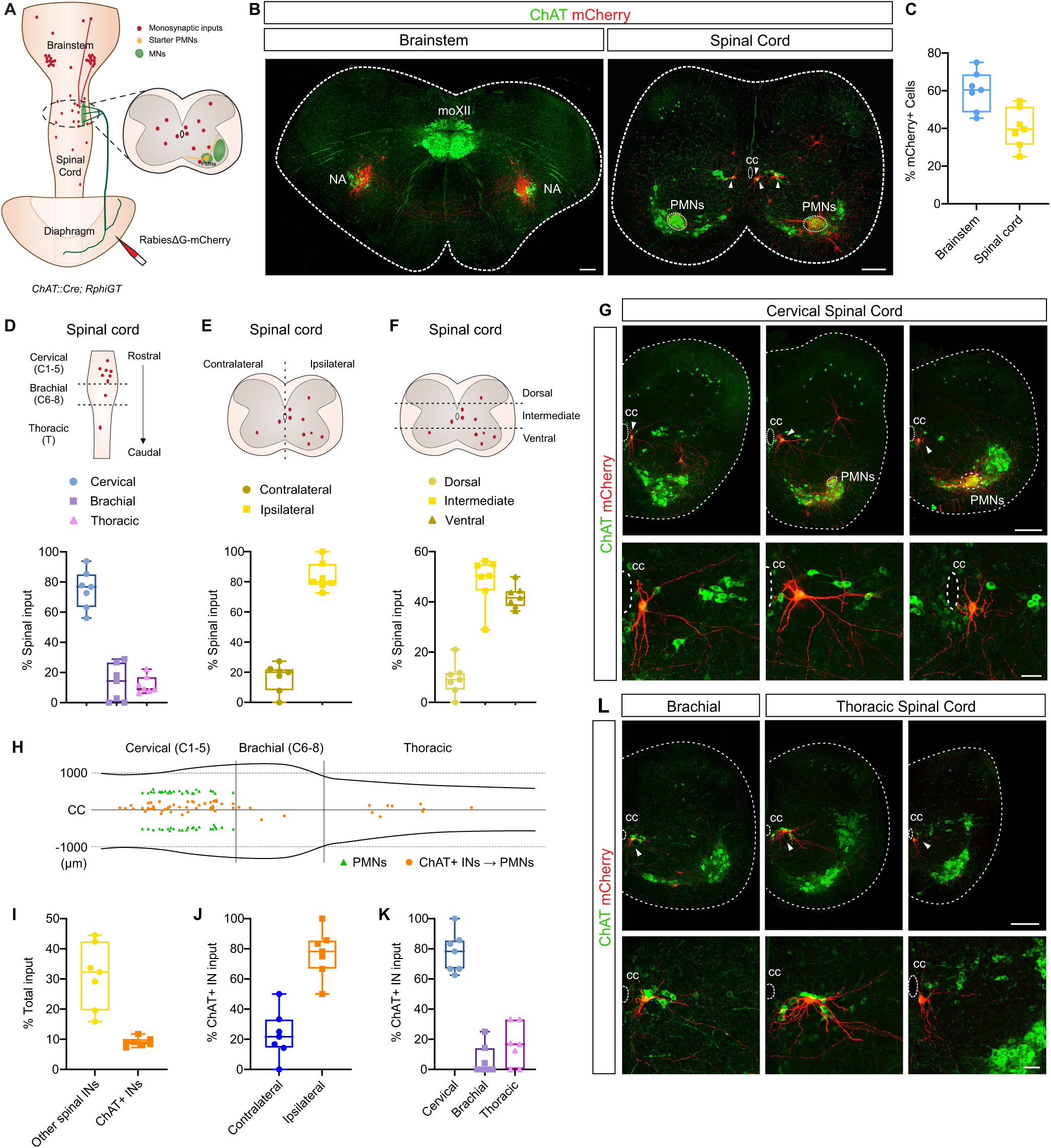
Distribution of monosynaptic inputs to phrenic motor neurons (PMNs) (A) Tracing strategy for mapping monosynaptic PMN inputs. (B) Examples of mCherry-labeled brainstem (rVRG) and spinal cord interneurons projecting to PMNs. moXII: hypoglossal motor nucleus, NA: Nucleus Ambiguus, cc: central canal. Scale bar = 200 μm. (C) Distribution of PMN inputs in the brainstem and spinal cord (n = 7). (D) Quantification of the rostrocaudal distribution of spinal inputs to PMNs. (E) Spinal cord PMN inputs are largely localized to the ipsilateral side. (F) Dorsoventral distribution of PMN inputs in the spinal cord. (G) Cholinergic interneurons (ChAT+ INs) around the central canal (cc) in the cervical spinal cord directly project to PMNs. Scale bar = 200 μm (top) and 50 μm (bottom). (H) Rostrocaudal distribution of ChAT+ INs projecting to PMNs (ChAT+ INs → PMNs). (I) ChAT+ INs comprise ∼10% of total PMN inputs. (J) Quantification of contralateral and ipsilateral ChAT+ INs projecting to PMNs. (K) Quantification of the rostrocaudal distribution of ChAT+ INs projecting to PMNs. (L) ChAT+ INs from the brachial and thoracic spinal cord directly project to PMNs. Scale bar = 200 μm (top) and 50 μm (bottom).

Next, we quantified the distribution of direct PMN inputs throughout the brain and spinal cord. All mCherry+ cells were located in either the brainstem or spinal cord and no mCherry+ cells were found in the cortex or cerebellum (data not shown). In agreement with a previous study^25^, we found that the majority (∼60%) of inputs to PMNs originated from the brainstem, mainly from the rostral ventral respiratory group (rVRG), consistent with rVRG being the major driver of PMN activation (Fig. 1B). In the brainstem, inputs were evenly distributed across the ipsilateral (to the injection site, 49.6%) and contralateral (50.4%) sides (Extended Data Fig. 2C). In addition, we also observed substantial (∼40%) PMN inputs originating from the spinal cord (Fig. 1B, C, G, and L, Extended Data Fig. 2F and G, Extended Data Video 1). Among spinal cord inputs, the large majority (∼80%) were located at cervical levels (C1-5). We also observed a small number of ascending interneurons from the brachial (C6-C8, ∼10%) and thoracic (∼10%) spinal cord (Fig. 1D). Most spinal cord input neurons (80%) were ipsilateral to the injection site (Fig. 1E) and distributed across the ventral (42.1%, Extended Data Fig. 2F), intermediate (48.5%), and dorsal (9.4%, Fig. 1F, Extended Data Fig. 2G) spinal cord. We calculated the PMN connectivity index (number of mCherry+ neurons/starter cell) and found that a single PMN can receive input from tens of neurons, ranging from a dozen to over a hundred. However, regardless of the number of total inputs one PMN receives, input neurons show a similar distribution throughout the brainstem and spinal cord (Extended Data Fig. 2D). Overall, we found that spinal interneurons are a major source of monosynaptic inputs to PMNs.

To further investigate the identity of spinal monosynaptic inputs to PMNs, we examined the neurotransmitter profile of mCherry+ interneurons. We found that cholinergic interneurons (ChAT+ INs), usually located around the central canal of the intermediate spinal cord (Fig. 1G and L), contributed around 10% of total inputs (25% of spinal inputs) to PMNs (Fig. 1I). The connectivity index for ChAT+ INs for individual injections was invariably ∼1/10^th^ of the total input, indicating that distinct neuronal populations provide a consistent proportion of PMN monosynaptic inputs (Extended Data Fig. 2D and E). While ChAT+ INs accounted for ∼50% of inputs from the intermediate spinal cord, inputs from both the dorsal and ventral spinal cord were derived exclusively from ChAT-INs (Extended Data Fig. 2F-H). Similar to the distribution of other spinal inputs, 77% of PMN-projecting ChAT+ INs were from the ipsilateral side (Fig. 1J), while 77.6% were located in the cervical C1-C5 spinal cord, largely overlapping with the rostrocaudal distribution of PMNs, and consistent with the overall distribution of total spinal inputs (Fig. 1H and K). We also observed a small subset of ascending ChAT+ INs at brachial and thoracic levels (Fig. 1H, K, and L), suggesting that these interneurons may mediate communication with other respiratory and non-respiratory MNs. Along the rostrocaudal extent of the spinal cord, ChAT+ INs accounted for 25%, 5% and 33% of inputs from the cervical, brachial and thoracic spinal cord, respectively (Extended Data Fig. 2I). Taken together, our rabies tracing experiments demonstrate that spinal ChAT+ INs provide significant input to PMNs.

### PMN-projecting spinal ChAT+ INs are morphologically and topographically distinct

Our tracing experiments revealed that ∼10% of PMN inputs correspond to a subset of ChAT+ INs in the cervical spinal cord (Fig. 1). This contrasts limb-innervating MNs (LMNs), which receive extensive input from multiple populations of excitatory and inhibitory spinal interneurons, with only about 2% of their inputs originating from ChAT+ INs^39,40^. This biased connectivity suggests that ChAT+ INs likely have important modulatory roles in respiratory behaviors. To investigate whether distinct spinal ChAT+ INs project to different MN subtypes, we injected RabiesΔG-mCherry virus into a representative limb muscle, the biceps, to label ChAT+ INs that project to LMNs and compared their distribution and morphology to PMN-projecting ChAT+ INs (Fig. 2A-D). To analyze the topographical distribution of ChAT+ INs, each mCherry+ ChAT+ IN was assigned a cartesian coordinate, with the midpoint of the spinal cord midline defined as (0,0). Interestingly, we found that PMN-projecting ChAT+ INs were located closer to the central canal on average, compared to LMN-projecting ChAT+ INs (Fig. 2E-F).

**Figure 2.**
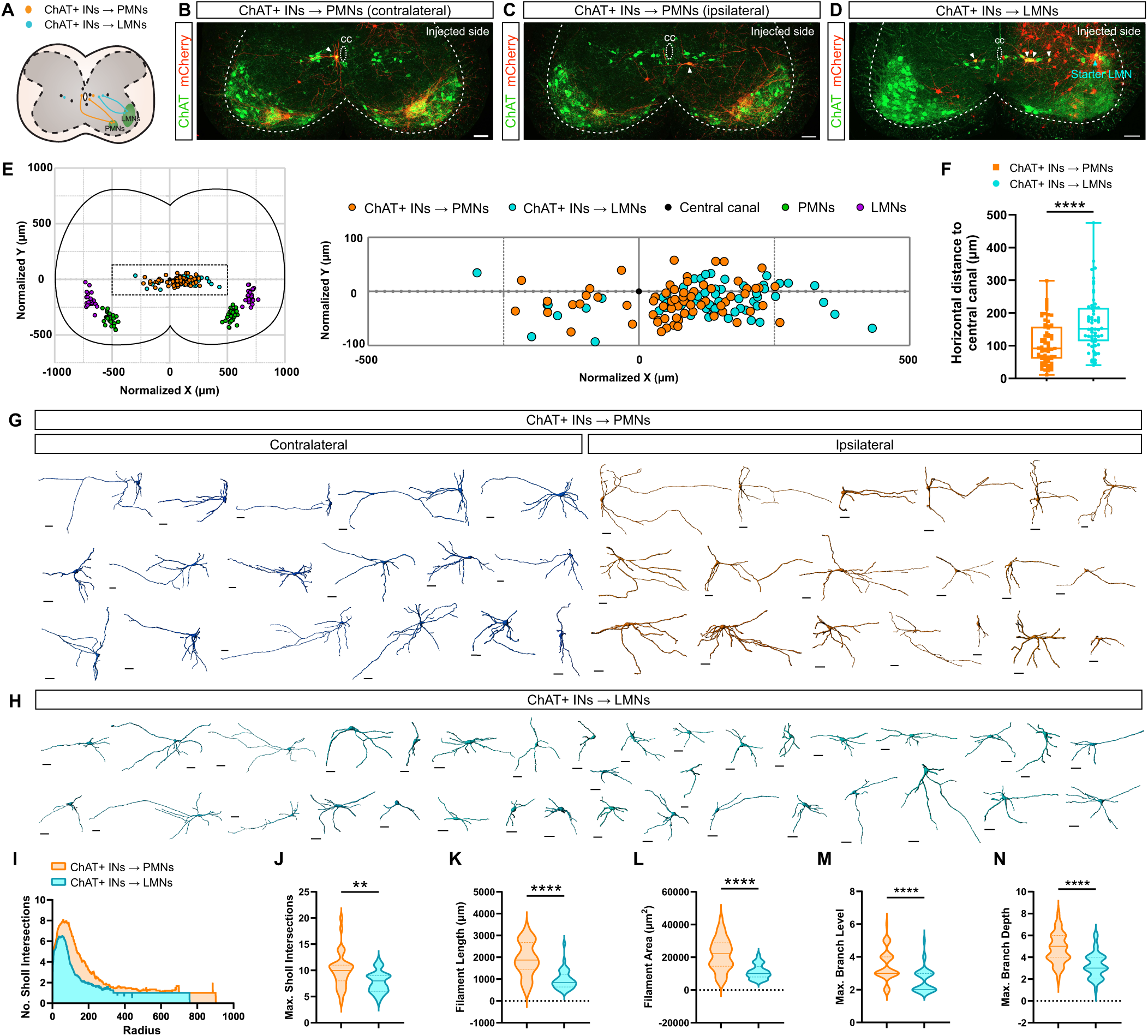
ChAT+ INs that project to PMNs are morphologically and topographically distinct. (A) Transsynaptic retrograde labeling of ChAT+ INs projecting to PMNs (ChAT+ INs → PMNs) and limb (biceps)-innervating MNs (ChAT+ INs → LMNs). (B-C) Representative images of contralateral (B) and ipsilateral (C) ChAT+ INs → PMNs. Scale bar = 100 μm. (D) Representative image of ChAT+ INs → LMNs. Scale bar = 100 μm. (E) Topographical distribution of ChAT+ INs → PMNs (n = 86 cells from 7 mice) and ChAT+ INs → LMNs (n = 69 cells from 5 mice). Rectangular region is enlarged to the right. (F) Quantification of ChAT+ INs → PMNs and ChAT+ INs → LMNs horizontal distance to the central canal. (G-H) Reconstruction of ChAT+ INs → PMNs (G) and ChAT+ INs → LMNs (H) morphology. Scale bar = 50 μm. (I) Sholl analysis of ChAT+ INs → PMNs (n = 36 cells) and ChAT+ INs → LMNs (n = 35 cells). (J-N) ChAT+ INs → PMNs have higher maximum Sholl intersections (J), greater overall dendritic length (K), cover a larger area (L), and have higher maximum branch level (M) and greater maximum branch depth (N) compared to ChAT+ INs → LMNs. ** p < 0.01, **** p < 0.0001

Next, we used Imaris software to reconstruct and examine the dendritic morphology of pre-motor ChAT+ INs by filament analysis (Extended Data Fig. 3A, Fig. 2G-H). First, to investigate whether there is diversity within PMN-projecting ChAT+ INs, we traced both contralateral and ipsilateral populations and compared their dendritic morphology. While all PMN-projecting ChAT+ INs had comparable dendritic length, dendritic area, and maximum dendritic branch level and depth, contralateral ones branched more at proximal dendritic levels and had a higher number of maximum Sholl intersections (Extended Data Fig. 3B-G). In addition, diverse dendritic orientation patterns were observed within PMN-projecting ChAT+ INs (Fig. 2G), suggesting that some morphological diversity exists even among ChAT+ INs targeting the same MN population.

We next compared the morphologies of ChAT+ INs projecting either to PMNs or LMNs. We found that PMN-projecting ChAT+ INs had more Sholl intersections, especially at proximal dendrites, and a higher number of maximum Sholl intersections (Fig. 2I-J). Moreover, ChAT+ INs targeting PMNs had a higher maximum branch level and depth, greater filament length (i.e. overall dendritic length), and covered a larger area (Fig. 2K-N), indicating that ChAT+ INs projecting to PMNs have a more complex dendritic morphology than ChAT+ INs targeting LMNs. Collectively, our findings suggest that ChAT+ INs that project to PMNs are topographically and morphologically distinct and may thus be integrated into respiratory, rather than locomotor, circuits.

### Pitx2+ V0_C_ neurons are the source of cholinergic synapses on PMNs

Large cholinergic synapses localized on the cell body and proximal dendrites of MNs are known as C-boutons^41^. To define whether cholinergic inputs on PMNs are analogous to the C-boutons observed on other MN subtypes, we investigated the distribution of cholinergic synapses on PMNs. Individual PMNs were traced by unilateral injection of RabiesΔG-mCherry virus into the diaphragm of control mice which lack G-protein expression, and therefore allowed RabiesΔG-mCherry virus to function as a retrograde tracer of only PMNs (i.e., no transsynaptic labeling). Cholinergic inputs on a single PMN were identified by immunostaining for vesicular Acetylcholine transporter (VAChT). We found that the majority of cholinergic synapses were located on the PMN soma (∼30%) and proximal dendrites (∼60%), with < 10% found on PMN distal dendrites (Extended Data Fig. 4), similar to the distribution of C-boutons on other MNs and consistent with ultrastructural studies^42^. The number of putative C-boutons on PMN cell bodies increased over time during early postnatal stages, but remained fairly unchanged from P4 to adulthood, indicating that PMNs receive consistent cholinergic input (Extended Data Fig. 5C).

Cholinergic V0 interneurons (V0_C_) that are derived from the Dbx1+ p0 progenitor domain and express Pitx2 post-mitotically, are the sole source of C-boutons on MNs^43^. To investigate whether cholinergic inputs to PMNs are also derived from V0_C_ interneurons, we genetically labeled Pitx2+ interneurons using *Pitx2::Cre; loxP-STOP-loxP-tdTomato* (*Pitx2^tdTom^*) mice. We found that over 90% of the immunoreactive Pitx2+ cells were tdTomato+ at e16.5, indicating robust recombination in *Pitx2^tdTom^* mice (Extended Data Fig. 5A-B). We labeled PMNs by intrapleural injection of cholera toxin subunit B (CTB) in adult *Pitx2^tdTom^* mice (Fig. 3A-B)^44^ and found that VAChT+ C-boutons on individual PMNs were colocalized with tdTomato (Fig. 3C-D), indicating that Pitx2+ V0_C_ interneurons are the source of cholinergic synapses on PMNs. To test whether all PMN C-bouton inputs are derived from Pitx2+ interneurons, we quantified the overlap between VAChT and tdTomato in *Pitx2^tdTom^* mice. We found that by P4 over 90% of C-boutons on PMN cell bodies were *Pitx2^tdTom^*+, and this was maintained into adulthood (P60) (Fig. 3E-N, Extended Data Fig. 5D). Collectively, our data show that Pitx2+ V0_C_ neurons are the source of PMN C-boutons largely localized on cell bodies and proximal dendrites.

**Figure 3.**
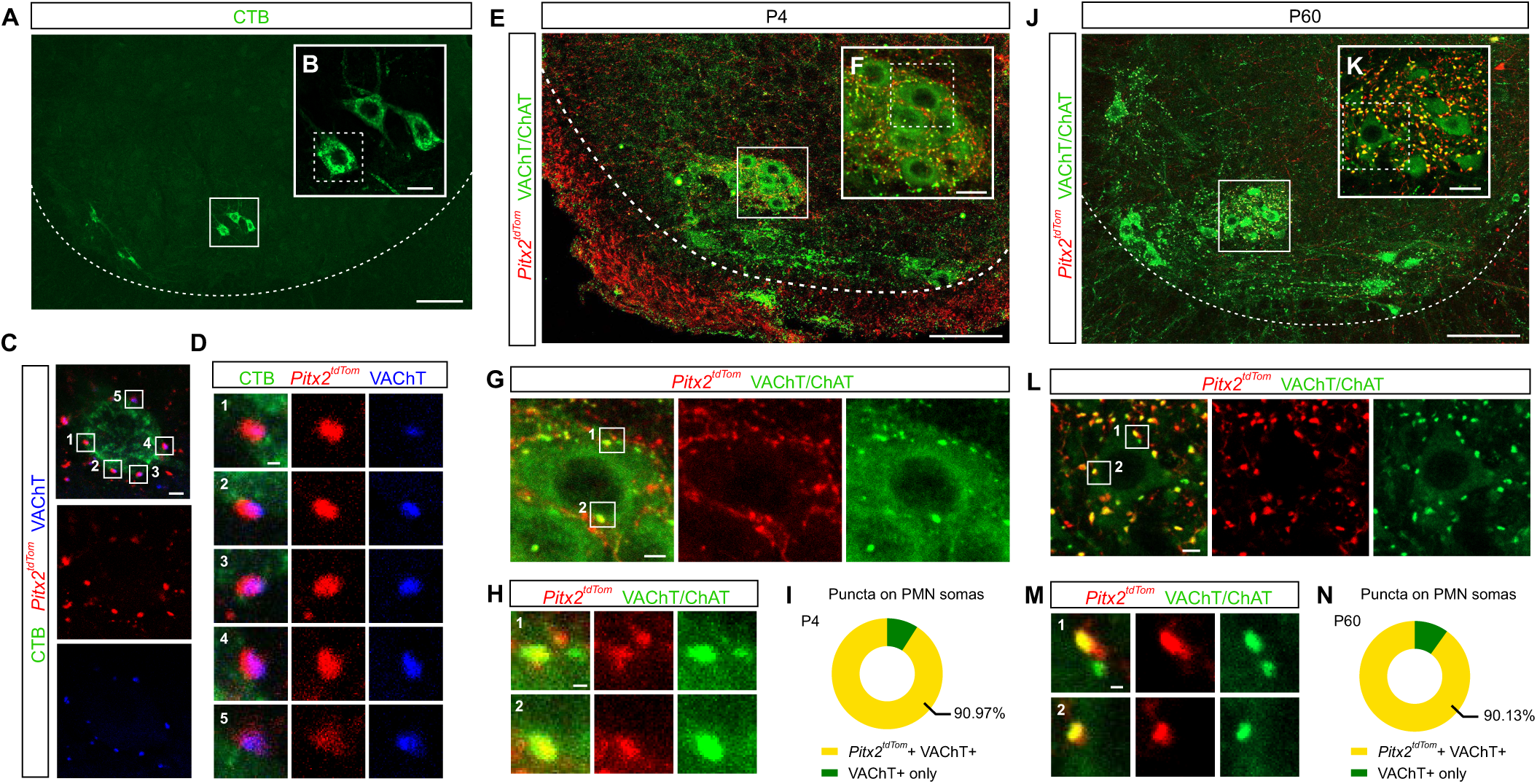
Pitx2-derived cholinergic synapses on PMNs. (A) Transverse section of the cervical spinal cord showing retrograde CTB labeling (green) in PMNs (squared region). Scale bar = 100 µm. (B) Enlargement of CTB-labeled PMNs (green) shown in (A). Square region indicates a PMN shown in (C). Scale bar = 20 µm. (C) Enlargement of a single CTB-labeled PMN (CTB, green) shown in (B) from a *Pitx2^tdTom^* adult mouse with synapses derived from *Pitx2^Tdtom^+* interneurons (red) and cholinergic synapses (VAChT, blue). Numbers indicate labeled synapses on the PMN soma. Scale bar = 5 µm (D) Magnification of numbered synapses from (C) showing colocalization of tdTomato and VAChT. Scale bar = 1 µm. (E-F) Representative images of VAChT+ puncta on PMNs in P4 *Pitx2^tdTom^* mice. Square region in (E) is enlarged in (F). Scale bar = 100 μm (E) and 20 μm (F). (G) Enlargement of the square region in (F). Numbers indicate examples of synapses on the PMN soma. Scale bar = 5 µm. (H) Magnification of numbered synapses from (G). Scale bar = 1 µm. (I) Percentage of VAChT+ puncta that are *Pitx2^tdTom^*+ on PMN somas at P4. (J-K) Representative images of VAChT+ puncta on PMNs in adult (P60) *Pitx2^tdTom^* mice. Square region in (J) is enlarged in (K). Scale bar = 100 μm (J) and 20 μm (K). (L) Enlargement of the square region in (K). Numbers indicate examples of synapses on the PMN soma. Scale bar = 5 µm. (M) Magnification of numbered synapses from (L). Scale bar = 1 µm. (N) Percentage of VAChT+ puncta that are *Pitx2^tdTom^*+ on PMN somas at P60.

### A subset of cervical Pitx2+ interneurons are integrated within respiratory circuits

Having revealed that a subset of Pitx2+ V0_C_ neurons near the central canal is anatomically connected to PMNs, we next addressed whether Pitx2+ interneurons are functionally integrated within respiratory circuits. This was first investigated by assessing whether their synaptic inputs and action potential output are correlated with respiratory network activity. We performed whole-cell patch clamp recordings from Pitx2+ interneurons located near the central canal, identified by tdTomato expression in *Pitx2^tdTom^* mice, in combination with extracellular recordings of C3/4 ventral roots in mid-sagittal-hemisected brainstem spinal cord preparations obtained from neonatal (P3-4) *Pitx2^tdTom^*mice (n = 5 preparations; Fig. 4A). Analysis of synaptic inputs recorded in voltage-clamp mode demonstrated that a subset of Pitx2+ interneurons within C3-4 (50%, n = 8, Fig. 4B) receive synaptic inputs (Frequency = 43 ± 21 Hz; Amplitude = 122 ± 142 pA) that are phase-locked with the respiratory motor output recorded from ventral roots (Fig. 4C; phase = 2.25 ± 0.9 degrees). There was no difference in the passive properties of Pitx2+ interneurons that received respiratory inputs compared to those that did not (Extended Data Table 1). Current-clamp recordings revealed that respiratory-related Pitx2+ interneurons exhibited either a depolarization of the membrane potential (n = 2; Amplitude = 7.14 mV) or bursts of action potential firing (n = 2, Frequency = 9.7 Hz) that were also phase-locked with respiratory-related ventral root output (Fig. 4D; Phase: 2.6 ± 0.7 degrees). These results indicate that at least a proportion of Pitx2+ interneurons located within the C3-4 cervical segments receive respiratory-related inputs and produce respiratory-related output. Interestingly, we did not find any respiratory-related Pitx2+ interneurons in more caudal cervical segments (C5-7, n = 10, Fig. 4B), suggesting that Pitx2+ interneurons in the region of PMNs are more likely to be integrated within respiratory circuitry, consistent with the distribution of PMN-projecting ChAT+ INs (Fig. 1H). Overall, these results confirm that spinal Pitx2+ interneurons are not only anatomically connected to PMNs but also functionally integrated within respiratory circuits.

**Figure 4.**
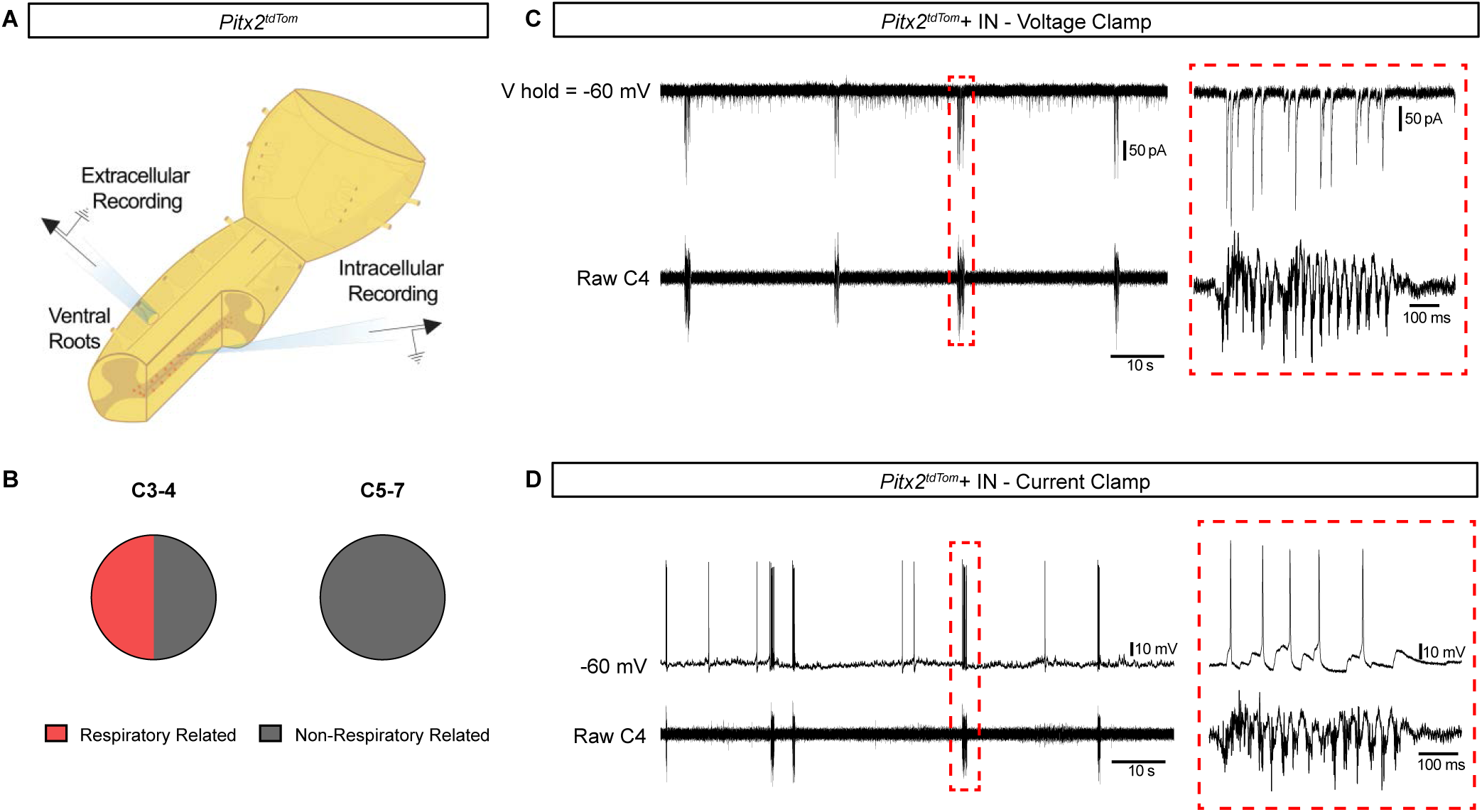
A subset of cervical Pitx2+ interneurons are integrated within respiratory circuits. (A) Schematic of the experimental setup showing extracellular ventral root recordings and intracellular whole-cell patch clamp recordings from individual *Pitx2^tdTom^*+ interneurons (red) in hemisected brainstem-spinal cord preparations from *Pitx2^tdTom^* neonatal mice. (B) Pie charts showing the relative proportion of respiratory-related (red) and non-respiratory-related (gray) Pitx2+ interneurons within and below the C3-4 spinal segments. (C) Example trace of voltage-clamp recording from a respiratory-related Pitx2+ interneuron (top) during ongoing respiratory burst (bottom). Red dotted box showing zoomed-in traces during a respiratory burst. (D) as in (C) but regarding current-clamp recording.

### Spinal ChAT+ INs modulate respiratory motor output

We next assessed the role of cervical Pitx2+ interneurons in modulating respiratory-related PMN output. This was achieved by pharmacologically blocking transmission at C-bouton synapses and measuring any subsequent effects on respiratory network output recorded from the C3/4 ventral roots of isolated brainstem spinal cord preparations obtained from neonatal (P2-4) mice (Fig. 5A). We utilized the M2 muscarinic receptor antagonist, methoctramine, because M2 receptors are the primary postsynaptic target for acetylcholine released at C-bouton synapses^45^. In line with C-boutons playing a role in facilitating motor output, we found that blocking M2 receptors with methoctramine (10 µM; n = 7 preparations; Fig. 5B) reversibly reduced the amplitude of respiratory-related activity (19 ± 9 % reduction, p = 0.0100; Fig. 5C). This reduction in amplitude was paralleled by an increase in the frequency of bursting (55 ± 34 % increase, p = 0.0107; Fig. 5D).

**Figure 5.**
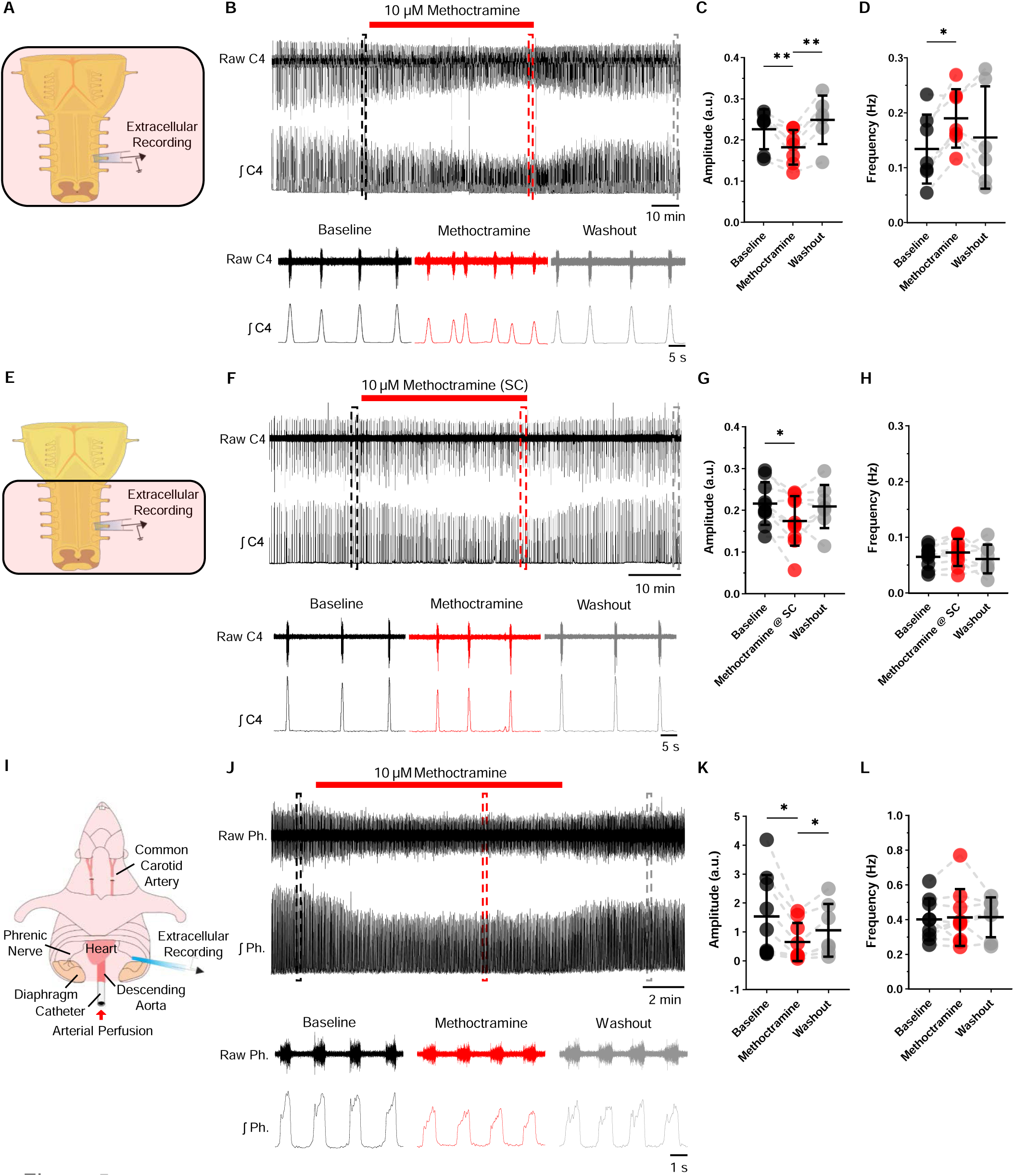
Effect of methoctramine on the respiratory motor output. (A) Brainstem-spinal cord preparation from neonatal mice. (B) Raw (top) and integrated/rectified (bottom) traces from the C4 ventral root during baseline, methoctramine (10 µM) and washout. a-c boxes indicate 40 seconds of recording at the end of each condition (black = baseline; red = methoctramine; grey = washout), expanded at the bottom. (C) Average respiratory-burst amplitude over the last 10 minutes during baseline (black), 10 µM methoctramine (red), and washout (grey); black lines show mean and SD (n = 7). (D) as in (C) but showing respiratory-burst frequency. (E) Experimental design to block M2 receptors at spinal levels only in the brainstem-spinal cord preparation from neonatal mice. (F) as in (B) but showing the effect of methoctramine at spinal level only. (G) and (H) as in (C) and (D) but showing the effect of methoctramine at spinal level only (n = 10). (I) Working heart-brainstem preparation from adult rats. (J) as (B) and (F) but showing the phrenic neurogram trace from the working heart-brainstem preparation. Square boxes indicate 7 seconds of recording at the end of each condition (black = baseline; red = methoctramine; grey = washout), expanded at the bottom. (K) and (L) as in (C) and (D) but showing the effects of methoctramine in the adult preparation. Data analyzed with mixed-effect model and Holm-Šídák’s multiple comparisons test. *p < 0.05, **p < 0.001

Since our experiments involved blockade of M2 receptors throughout the brainstem and spinal cord, we next wanted to delineate the specific contribution of spinal cord interneurons to cholinergic modulation of respiratory motor output. This was achieved by using a “split bath” preparation in which the brainstem and spinal cord compartments were perfused separately (Fig. 5E). This enabled us to block M2 receptors in the spinal cord only. We hypothesized that the reduction in the amplitude of respiratory-related output when methoctramine was applied to whole preparations could be explained, at least in part, by blockade of M2 receptors at C-bouton synapses on PMNs. In line with our hypothesis, application of methoctramine (10 µM; n =10 preparations; Fig. 5F) to the spinal compartment of split bath preparations only led to a reversible reduction in the amplitude of respiratory-related output recorded from ventral roots (20 ± 18 % reduction, p = 0.0120; Fig. 5G), with no change in the frequency of bursting (12 ± 14 % increase, p = 0.1221; Fig. 5H).

Given that our in vitro experiments relied on isolated brainstem spinal cord preparations that are only viable when obtained from neonatal animals, we next extended our analysis to the working heart-brainstem preparation, which can be obtained from adult rodents (Fig. 5I). In preparations obtained from adult rats, we recorded respiratory-related output from the phrenic nerve with extracellular electrodes whilst methoctramine (10 µM, n = 9 preparations; Fig. 5J) was applied to the perfusate. Consistent with our recordings from neonatal tissue, we again found that methoctramine caused a reversible reduction in the amplitude of respiratory-related output recorded from the phrenic nerve (57 ± 15 % reduction; p = 0.0294; Fig. 5K). We did not observe any change in the frequency of phrenic nerve discharge upon methoctramine administration (2 ± 16 % increase; p = 0.6432; Fig. 5L). Interestingly, we found that methoctramine also elicited a reduction in the amplitude of respiratory-related activity recorded from external intercostal muscles via electromyography (44 ± 16 % reduction; p = 0.007, data not shown).

Taken together, these results confirm the existence of a spinal cholinergic pathway, acting via M2 receptors, that modulates the amplitude of PMN output in both neonatal and adult mice. These data are consistent with modulation of respiratory motor output by local Pitx2+ interneurons and their C-bouton contacts with PMNs.

### Spinal cholinergic interneurons are activated under hypercapnia

Given the functional integration of Pitx2+ interneurons into respiratory circuits and their ability to alter PMN output in both in vitro neonatal isolated brainstem-spinal cord preparations and in situ adult working heart-brainstem preparations, we next investigated whether Pitx2+ V0_C_ neuron activity can modulate respiratory behaviors in vivo. In order to visualize V0_C_ activation, we utilized *ChAT::eGFP* mice, in which both ChAT+ INs and PMNs are labeled by GFP, but can be distinguished by their locations. While PMNs are clustered and located in the ventral horn of the cervical spinal cord, ChAT+ INs are scattered throughout the spinal cord, with the distinct PMN-projecting Pitx2+ V0_C_ subset localized near the central canal (Fig. 6A).

**Figure 6.**
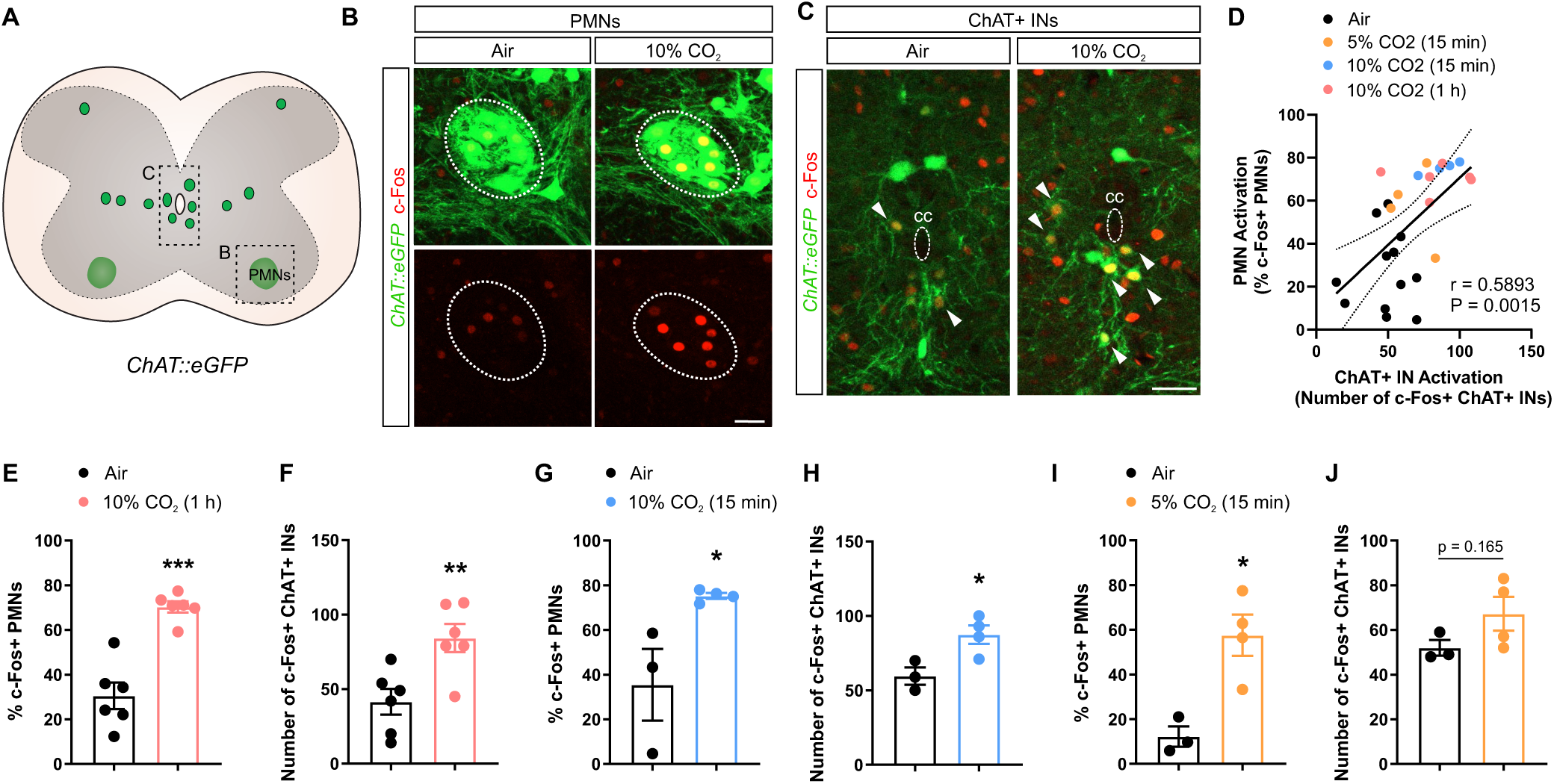
ChAT+ INs are highly activated under a hypercapnic gas challenge. (A) ChAT+ MNs and INs are labeled by GFP in *ChAT::eGFP* mice. Regions including PMNs and ChAT+ INs around the central canal are shown in (B) and (C), respectively. (B-C) Both PMNs (B) and ChAT+ INs (C) are highly activated, as indicated by high c-Fos expression (red), after exposure to 10% CO_2_. White arrows indicate c-Fos+ ChAT+ INs. Scale bar = 40 μm. (D) PMN activation is positively correlated to ChAT+ IN activation. (E, G, and I) Percentage of c-Fos+ PMNs after 10% CO_2_ for 1 hour (E), 10% CO_2_ for 15 minutes (G), and 5% CO_2_ for 15 minutes (I). (F, H, and J) Number of c-Fos+ ChAT INs after 10% CO_2_ for 1 hour (F), 10% CO_2_ for 15 minutes (H), and 5% CO_2_ for 15 minutes (J). * p < 0.05, ** p < 0.01, *** p < 0.001

In locomotor circuits, Pitx2+ V0_C_ interneurons increase MN excitability to ensure that a sufficient motor output is generated during demanding tasks, such as swimming^43,46^. We therefore hypothesized that cervical V0_C_ interneurons may contribute to increasing respiratory output during environmental or behavioral conditions that are associated with increased metabolic demand. To measure neuronal activation under a respiratory challenge, we exposed *ChAT::eGFP* mice to hypercapnic conditions in a plethysmography chamber to simulate increased metabolic demand. Under atmospheric air conditions (79% N_2_, 21% O_2_), only 30% of PMNs were activated, as indicated by the expression of the early immediate gene c-Fos. In contrast, more than 60% of PMNs were activated under either intense (10% CO_2_ for either 1 hr or 15 min) or moderate (5% CO_2_ for 15 min) hypercapnic conditions (Fig. 6B, E, G and I). Moreover, the c-Fos mean intensity of PMNs was significantly increased after exposure to 10% CO_2_ (Extended Data Fig. 6B-C), indicating that both the number of recruited PMNs and single PMN activity are increased under an intense hypercapnic challenge, as previously described^47,48^. CO_2_ exposure does not lead to broad non-specific activation of spinal cord neurons, as we found that the number of c-Fos+ cells at cervical levels of the spinal cord, excluding MNs, was comparable to baseline levels (i.e. air exposure) after a 10% CO_2_ hypercapnic challenge for 1 hour (Extended Data Fig. 6A).

Since PMNs are highly activated under hypercapnia, a hypercapnic gas challenge can serve as a powerful paradigm to examine whether spinal interneurons modulate PMN activation under an environmental challenge. We found that ChAT+ INs were highly activated under an intense (15 min or 1 hr of 10% CO_2_; Fig. 6C, F, and H), but not under a moderate (15 min of 5% CO_2_) hypercapnic challenge (Fig. 6J), suggesting that ChAT+ IN activation and recruitment is dependent on the intensity level of the stimulus. c-Fos+ ChAT+ INs were located in close proximity to the central canal at cervical levels of the spinal cord, and thus are likely to directly project to PMNs and correspond to the Pitx2+ interneurons with respiratory-related activity in our in vitro recordings. While the number of activated ChAT+ INs recruited under an intense hypercapnic challenge increased, the c-Fos mean intensity of ChAT+ INs did not change significantly, unlike PMNs (Extended Data Fig. 6B-E), suggesting that recruitment of additional ChAT+ INs may contribute to increased PMN activation. Consistent with this idea, we found that the number of activated ChAT+ INs was positively correlated to the percentage of activated PMNs (Fig. 6D).

### Cholinergic interneuron silencing impairs the response to hypercapnia

Next, we set out to determine the contribution of ChAT+ INs to increased breathing in response to hypercapnia. To address this, we utilized a 2-chamber whole body plethysmography system to measure the breathing patterns in mice in which cholinergic neurotransmission has been removed from spinal ChAT+ INs using Cre/lox genetic strategies (Extended Data Fig. 7A). We initially utilized *ChAT^flox/flox^*; *Dbx1::Cre* (*Dbx1^ΔChAT^*) mice to ensure early and efficient *ChAT* deletion, as *Dbx1::Cre* targets p0 progenitors that give rise to Pitx2+ V0_C_ interneurons. We validated *Dbx1^ΔChAT^*mice by examining the expression of ChAT in VAChT+ puncta on PMNs in adult (P120) mice. We found that over 95% of VAChT+ terminals on PMNs were devoid of ChAT expression in *Dbx1^ΔChAT^* mice, indicating that cholinergic transmission from V0_C_ neurons to PMNs was largely eliminated in these mice (Extended Data Fig. 7B-D). We did not observe any changes in the number of VAChT+ synapses on PMN somas in adult *Dbx1^ΔChAT^*mice (Extended Data Fig. 7E).

We then performed plethysmography experiments in adult *Dbx1^ΔChAT^*mice and their paired control littermates (*ChAT^flox/flox^* or *ChAT^flox/+^*). Each pair of mice was subjected to 45 minutes of normal air (79% N_2_, 21% O_2_) followed by 15 minutes of either 10% CO_2_ (10% CO_2_, 69% N_2_, 21% O_2_) or 5% CO_2_ (5% CO_2_, 74% N_2_, 21% O_2_) (Fig. 7A). When exposed to a hypercapnia challenge, control mice increase both their respiratory frequency and depth. Consistent with this, we found a significant increase in both frequency and tidal volume (the amount of air inhaled during a normal breath), resulting in a ∼3-fold increase in minute ventilation (the volume of air inhaled per minute) in control mice after exposure to a 10% CO_2_ hypercapnic challenge (Extended Data Fig. 8A-H). When comparing *Dbx1^ΔChAT^* mice and their control littermates, we found that they exhibited similar breathing behaviors under normal air conditions. All breathing parameters, including frequency, minute ventilation, tidal volume, and peak inspiratory and expiratory flow showed similar distribution in *Dbx1^ΔChAT^* and control mice (Extended Data Fig. 8C, E, G, I, K). However, after exposure to a 10% CO_2_ hypercapnic challenge, the distribution peaks for all respiratory parameters, excluding frequency, were shifted to a lower value in *Dbx1^ΔChAT^* mice, indicating that these mice had a compromised response to hypercapnia (Extended Data Fig. 8D, F, H, J, L). After normalization to their paired control littermates, *Dbx1^ΔChAT^* mice had significantly decreased tidal volume (∼11% reduction, p = 0.0291), minute ventilation (∼13% reduction, p = 0.0150), peak inspiratory flow (PIF, ∼15% reduction, p = 0.0035) and peak expiratory flow (PEF, ∼11% reduction, p = 0.0281) compared to control mice during the hypercapnia challenge (Extended Data Fig. 8F, H, J, L), consistent with the altered distribution of respiratory parameters we observed.

**Figure 7.**
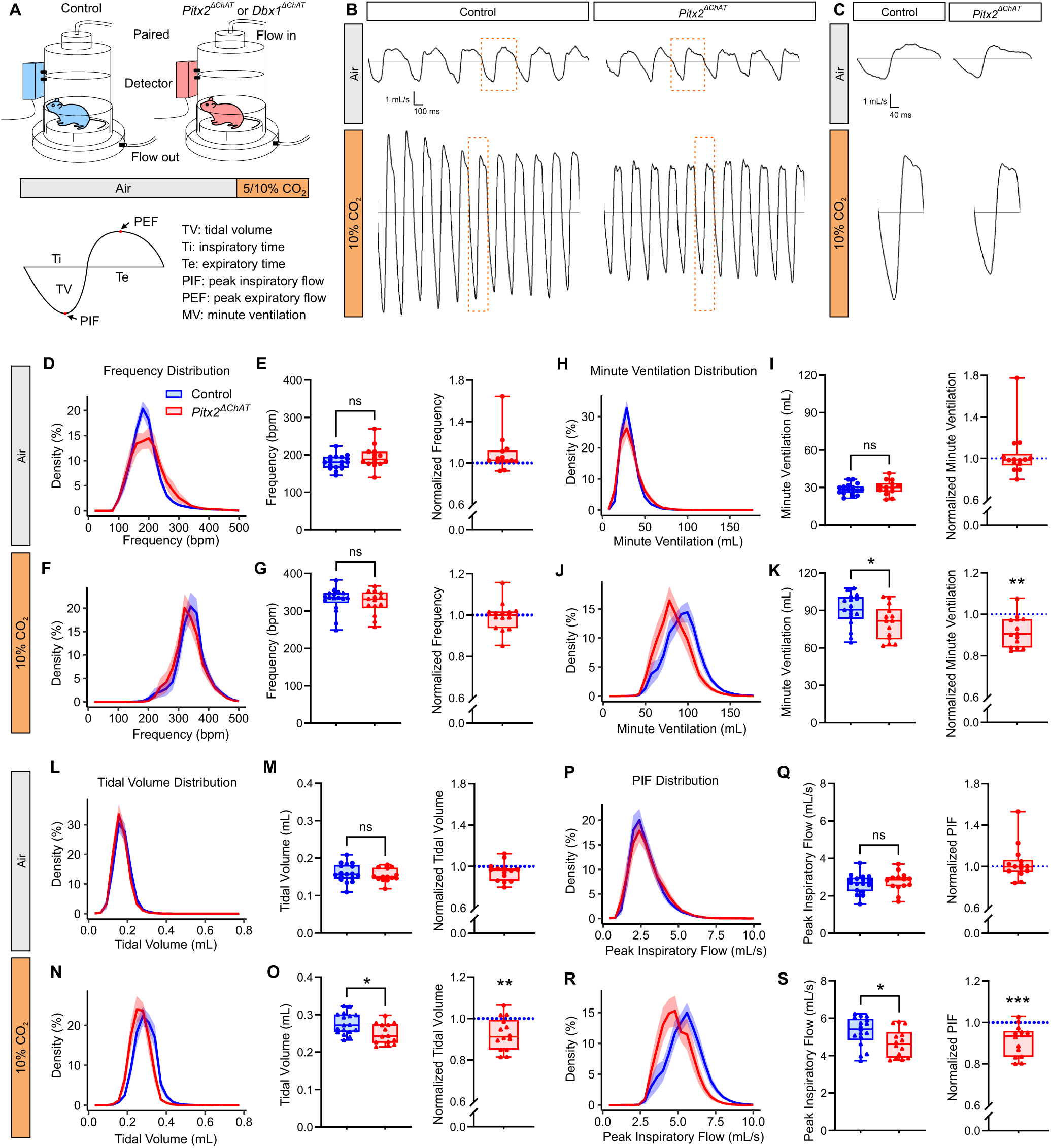
Cholinergic interneuron silencing impairs the response to hypercapnia. (A) Experimental design for whole body plethysmography and schematic of a representative breath. A *Pitx2^ΔChAT^* or *Dbx1^ΔChAT^* mouse was paired with a sex-matched control littermate and the mice were exposed to normal air conditions (21% O_2_, 79% N_2_) for 45 minutes, followed by 15 minutes of hypercapnia (5% CO_2_, 21% O_2_, 74% N_2_ or 10% CO_2_, 21% O_2_, 69% N_2_). (B-C) Examples of breath traces under normal air and 10% CO_2_ in *Pitx2^ΔChAT^* mice and their control littermates. A single breath was enlarged in (C). (D, H, L, and P) Breath frequency (D), minute ventilation (H), tidal volume (L), and PIF (P) distribution under normal air in *Pitx2^ΔChAT^*and control mice (n = 14-17 per group). (E, I, M, and Q) Mean and normalized frequency (E), minute ventilation (I), tidal volume (M), and PIF (Q) under normal air in *Pitx2^ΔChAT^*and control mice. (F, J, N, and R) Breath frequency (F), minute ventilation (J), tidal volume (N), and PIF (R) distribution under 10% CO_2_ in *Pitx2^ΔChAT^*and control mice (n = 14-17 per group). (G, K, O, and S) Mean and normalized frequency (G), minute ventilation (K), tidal volume (O), and PIF (S) under 10% CO_2_ in *Pitx2^ΔChAT^* and control mice.

Since Dbx1 is broadly expressed at early embryonic stages, we wanted to further restrict our manipulations to V0_C_ postmitotic interneurons. Therefore, we repeated our plethysmography experiments in *ChAT^flox/flox^*; *Pitx2::Cre* (*Pitx2^ΔChAT^*) mice, which restricted *ChAT* deletion to Pitx2+ interneurons. Similar to *Dbx1^ΔChAT^*mice, over 95% of VAChT+ terminals on PMNs did not express ChAT in adult *Pitx2^ΔChAT^* mice, indicating efficient ChAT deletion from V0_C_ neurons (Extended Data Fig. 7F-H). We did not observe a decrease in VAChT+ terminals on PMNs in these mice (Extended Data Fig. 7I). We exposed *ChAT^Pitx2Δ^* mice and their control littermates to both normal air and hypercapnic conditions (Fig. 7A). Since we found that the activation of ChAT+ INs was intensity-dependent (Fig. 6), we first investigated how blocking cholinergic neurotransmission from V0_C_ interneurons might impact breathing under a moderate hypercapnic challenge. Interestingly, when we exposed *Pitx2^ΔChAT^* mice to either normal air or 5% CO_2_, their breathing patterns were indistinguishable from their control littermates, suggesting that PMN cholinergic modulation might be increasingly important under more intense hypercapnic challenges (Fig. 7D-E, H-I, L-M, P-Q, Extended Data Fig. 9).

Consistent with our observations in *Dbx1^ΔChAT^* mice, the tidal volume, minute ventilation, PIF, and PEF distributions were all shifted toward lower values (i.e. more breaths having a lower value) under 10% CO_2_ hypercapnia, but not under normal air conditions, in *Pitx2^ΔChAT^* mice (Fig. 7B-D, F, H, J, L, N, P, R, Extended Data Fig. 8M-N). In addition, the tidal volume (∼8% reduction, p = 0.0033), minute ventilation (∼8% reduction, p = 0.0011), PIF (∼9% reduction, p = 0.0007), and PEF (∼7% reduction, p = 0.0104), but not the breathing frequency (< 1%), were significantly decreased in *Pitx2^ΔChAT^*mice under 10% CO_2_ (Fig. 7E, G, I, K, M, O, Q, S, Extended Data Fig. 8N). Taken together, our findings indicate that cholinergic modulation of PMNs, through direct C-bouton contacts from spinal Pitx2+ V0_C_ interneurons, increases PMN output under a hypercapnia challenge. Collectively, our data reveal a novel function for spinal cholinergic interneurons in the adaptive control of breathing.

## Discussion

Phrenic motor neurons (PMNs) generate the final output of respiratory motor circuits and have traditionally been thought to eschew inputs from the spinal cord and act as executioners of brainstem motor commands. Here, we combined mouse genetics, rabies-mediated viral tracing, electrophysiology, and behavioral experiments to demonstrate a novel role for a subset of spinal cholinergic interneurons in the facilitation of PMN output and increase in tidal volume in response to hypercapnia. Our data suggest that, far from being a static relay station for brainstem motor commands, PMNs integrate a range of modulatory inputs to match motor output to environmental or metabolic demands. Below, we discuss our findings in the context of PMN modulation, respiratory circuits, spinal interneuron diversity, and potential roles in promoting recovery following spinal cord injury.

While spinal interneurons with projections to PMNs have been anatomically and electrophysiologically described in multiple species, the contribution of genetically-defined classes of spinal interneurons to distinct aspects of breathing remains unclear. Previous mapping of PMN inputs through transsynaptic viral approaches has revealed varying amounts of spinal interneuron inputs, ranging from substantial in adult rats to negligible in neonatal mice^20,23,25,26^. In addition to potential species differences, other potential sources of variability may be the tropism of the different viruses used (e.g. PRV vs Rabies) or the age of the injected animals (neonatal vs adult), suggesting temporally dynamic inputs to PMNs that may be developmentally gained or lost. In addition to anatomical studies, microelectrode and multielectrode array recordings from the cervical spinal cord have identified a number of interneurons with both inspiratory and expiratory related activity, indicating that complex spinal circuits may be involved in PMN modulation downstream of brainstem circuits^29,30,49–53^. However, unlike well-described functions for cardinal interneuron classes in locomotion, mapping respiratory functions to these populations has been elusive. For example, Renshaw cells that respond to PMN stimulation and are spontaneously active during inspiration have been identified but appear to be rare^54–56^. Ablation or inhibition of V2a neurons changes the breathing frequency, but these effects are mediated primarily through the brainstem rhythm-generating pre-Botzinger Complex and accessory respiratory muscles rather than PMN modulation^57,58^. We find that Pitx2+ interneurons at cervical levels of the spinal cord directly project to PMNs, produce respiratory-related output, and modulate breathing amplitude under an environmental challenge, providing both anatomical and functional evidence for a respiratory role for V0_C_ interneurons. To our knowledge, this is the first time a genetically-defined spinal cord interneuron population has been implicated in PMN modulation and the adaptive control of breathing in healthy animals.

What are the inputs to spinal respiratory cholinergic interneurons, and how do they fit into the broad respiratory network? Putative limb MN-projecting cholinergic interneurons in the lumbar spinal cord increase the excitability of MNs involved in locomotion to ensure robust firing and sufficient MN output during demanding tasks such as swimming^43,46,59^. These interneurons receive input from corticospinal neurons, locomotor central pattern generator circuits, descending serotonergic inputs and polysynaptic inputs from sensory afferents to adjust their activity^43,60^. While respiratory V0_C_ neurons may also receive synaptic inputs from these sources, their activation by elevated CO_2_ levels in the absence of locomotor activity suggests alternative or additional inputs. One possibility is that they receive inputs from chemosensitive areas in the brainstem such as the retrotrapezoid nucleus (RTN) or serotonergic neurons that stimulate breathing in response to elevated CO_2_^5,61^; in fact, we do observe serotonergic axons in close proximity to mCherry+ ChAT+ INs in our rabies tracing experiments. In addition, respiratory V0_C_ neurons may receive inputs from pre-phrenic respiratory areas such as the rVRG; since cholinergic interneurons augment PMN output, the two populations might share common inputs. Previous studies have detected rVRG axons around cervical pre-phrenic interneurons^23,62^, and our recordings, which show synaptic currents phase locked with respiratory output, support this hypothesis. Respiratory V0_C_ interneurons may also receive inputs from areas that modulate breathing in response to different arousal states, such as the locus coeruleus^63^. In addition to cholinergic Pitx2+ V0_C_ interneurons, we also identified a number of excitatory and inhibitory PMN-projecting interneurons (Extended Data Fig. 2F and G) mostly localized in the cervical spinal cord, although it is worth noting that these are much less numerous than spinal cord interneurons projecting to limb MNs (compare Fig. 2B-C to 2D)^39,40,64^. Mapping the inputs to cholinergic, and all spinal respiratory interneurons broadly, will begin to dissect the regulatory influence of distinct respiratory populations on PMN excitability and function and provide insights into the circuits that modulate and enhance breathing behaviors. Altogether, our data suggests a more prominent contribution of spinal cord interneurons to the regulation of breathing than previously suggested.

While c-Fos experiments have limitations in detecting activation patterns with fine temporal resolution, we find that the hypercapnia challenge increases both the number of c-Fos expressing PMNs and the intensity of c-Fos expression, presumably corresponding to increases in both MN recruitment and firing rate. Conversely, only the number of c-Fos expressing ChAT+ INs changed, suggesting that V0_C_ recruitment is the predominant modality for increasing PMN gain and diaphragm output, similar to the recruitment of V2a interneuron subtypes to generate high speed movements^65,66^. Both rabies virus tracing and C-bouton quantification at PMN cell bodies reveal that there might be some variability in the number of V0_C_ inputs to each PMN. Although we did not attempt to correlate C-bouton number to cell body size, one possibility is that large, fast-fatigable PMNs receive more C-bouton inputs. Fast MNs have been shown to have a higher density of C-boutons and we have previously found greater effects of muscarine on fast MN output^59,67,68^. In addition to the hypercapnia response, V0_C_ recruitment and PMN modulation might become increasingly important for augmenting PMN output during expulsive behaviors, like coughing and sneezing, and exercise, when even greater PMN activation is required^69^.

While comprising a relatively small subset of interneurons within the spinal cord, V0_C_ interneurons exhibit remarkable molecular and anatomical diversity, and extensively innervate spinal MNs. Despite their extensive projections to MNs, there seems to be some selectivity in their targeting. For example, ocular and cremaster MNs lack cholinergic inputs, while MNs innervating large proximal muscles receive greater numbers of inputs than those innervating small distal muscles^67,68,70,71^. In addition, postsynaptic clustering of certain ion channels differs among cholinergic synapses on different MN subtypes^72^. Within locomotor circuits, there are at least two populations of V0_C_ neurons, one of which projects exclusively to ipsilateral targets while another projects either contralaterally or bilaterally, indicating that distinct programs underlie their connectivity^39,43^. Recent single nucleus (sn)RNA-seq analysis of cholinergic interneurons in adult mice identified eight transcriptionally distinct cholinergic interneuron clusters, two of which expressed Pitx2, further supporting the idea that there is significant diversity within this interneuron population, despite their common developmental origin from Dbx1-expressing progenitors^73^. Therefore, it is likely that distinct cholinergic interneurons innervate specific MN subtypes to mediate unique functions. We now show that V0_C_ interneurons participate in diverse behavioral responses, modulating MN output to control limb movements as well as breathing.

While our data clearly point to the existence of a subset of respiratory-related V0_C_ neurons, we cannot definitively determine whether these interneurons exclusively project to PMNs or also target limb MNs. Dually-projecting cholinergic interneurons have been observed in adult rats^74^, and even within the phrenic-projecting V0_C_ population we observe considerable diversity in dendritic orientation and rostrocaudal distribution, indicating that smaller subdivisions, with distinct inputs and outputs, might exist even within this subset. For example, V0_C_ neurons at thoracic levels may project to both phrenic and intercostal MNs to coordinately increase respiratory output. Interestingly, we found that in adult perfused preparations, methoctramine also elicited a reduction in the amplitude of respiratory-related activity in external intercostal muscles, although we did not identify the source of this modulation. Mapping the synaptic inputs and outputs of individual V0_C_ neurons will reveal the extent of common V0_C_ inputs onto different MNs.

Despite well-defined morphogen gradients and transcriptional programs that specify cardinal classes of interneurons, how these major classes are subdivided into smaller, distinct subsets with discrete projections and functions is just beginning to emerge^75,76^. The heavily biased cervical-level distribution of respiratory V0_C_ interneurons suggests that perhaps genetic programs involved in rostrocaudal patterning might shape their identity. For example, Hox transcription factors control MN diversity along the rostrocaudal axis of the spinal cord and play critical roles in the specification of MN subtypes, including PMNs^77–79^. Recent evidence suggests that Hox proteins may also have broad roles in the specification and connectivity of spatially-segregated interneuron subtypes^80^. The integration of cholinergic interneurons into respiratory or locomotor circuits suggests that a developmental logic may dictate their selective targeting, both to specific MNs and from distinct pre-synaptic partners. Whether this selective connectivity is explicitly linked to early genetic programs that also define their topography and morphology, or shaped by activity-dependent mechanisms, remains to be determined.

While the role of spinal interneurons in the canonical control of breathing has remained somewhat elusive, recent studies have highlighted their critical function in respiratory plasticity and recovery after spinal cord injury (SCI). For instance, animal studies have demonstrated that changes in the function or connectivity of propriospinal neurons are crucial for improving breathing function during both acute and chronic stages of injury^31–33,36,37^. Excitatory interneurons are particularly important for recovery, as evidenced by models of non-traumatic cervical myelopathy^32^ and C2 hemisection SCI^31^. Specifically, in C2 hemisection SCI, increased connectivity between V2a interneurons and PMNs has been linked to the spontaneous recovery of breathing^81,82^. Activation of V2a interneurons can restore function in the paralyzed hemidiaphragm following a C2 hemisection injury, while silencing these neurons significantly hinders recovery^33^. Although the role of ChAT+ INs in recovering breathing function after SCI remains to be elucidated, their importance in neurodegenerative conditions is suggested by studies of V0_C_ neurons in locomotor circuits in ALS^83–85^. Given our findings supporting a role for V0_C_ interneurons in facilitating diaphragm activation during hypercapnia, they represent a promising neuronal population that can be coopted to counteract the reduced hypercapnic drive response seen in cervical SCI patients^86^ and enhance recovery after injury.

## Methods

### Animals

*RphiGT* (JAX# 024708)^38^, *ROSA26Sor^tm9(CAG-tdTomato)^* (Ai9, JAX# 007909)^87^, *ChAT::eGFP* (JAX# 030250)^88^, *ChAT^flox/flox^* (JAX# 016920)^89^, *ChAT::Cre*^90^*, Dbx1::Cre*^91^, and *Pitx2::Cre*^92^ lines were generated as previously described and maintained on a mixed background. No more than five adult mice were housed in a microisolator cage at one time under a 12-hr light/dark cycle. Procedures and mouse maintenance performed in the United States were executed in accordance with protocols approved by the Institutional Animal Care Use Committee of Case Western Reserve University (assurance number: A-3145-01, protocol number: 2015-0180). Procedures performed in the United Kingdom were conducted in accordance with the UK Animals (Scientific Procedures) Act 1986, were approved by the University of St Andrews Animal Welfare Ethics Committee and were covered under Project Licences (P6F7B721E and PP8253850) approved by the Home Office. Experiments carried out in Canada were approved by the Animal Care Committee at the University of Calgary (AC19-0037).

### Rabies-based monosynaptic tracing

RabiesΔG-mCherry virus production and monosynaptic tracing were performed as previously described^93,94^. Briefly, rabies injection solution was made by mixing RabiesΔG-mCherry virus (titer of around 1^10^ TU/ml) with silk fibroin (Sigma# 5154)^95^ at a 2:1 ratio. 1-1.5 μl of rabies injection solution was unilaterally injected into the diaphragm or biceps muscle of *ChAT::Cre; RphiGT* mice at P4 using a nano-injector (Drummond). Mice were sacrificed 7 days post-injection (P11). Specificity of the unilateral injection of the diaphragm/PMN infection was confirmed by checking the mCherry fluorescent signal in the diaphragm and spinal cord (from cervical to lumbar levels, Extended Data Fig.1C). Fluorescent signal at the ventral roots was used as an indicator of starter MN labeling. No signal at ventral roots outside the C3-C5 spinal cord was detected in animals with phrenic-specific labeling. 100 μm-thick consecutive sections from the cerebrum to the spinal cord were harvested by Leica VT1000S vibratome. Sectioning was stopped if no mCherry+ cells were observed in more than 20 consecutive sections (2 mm). The connectivity index was calculated by dividing the number of mCherry+ labeled cells by the number of starter PMNs in individual animals.

### Immunohistochemistry

Mice aged older than P12 were anesthetized with either a Ketamine/Xylazine cocktail or Pentobarbital and underwent transcardial perfusion with ice-cold phosphate buffered saline (PBS; pH 7.4 without Ca^2+^ or Mg^2+^) to remove the blood, followed by ice-cold 4% paraformaldehyde (PFA). Neonatal mice were dissected acutely after being anesthetized with Ketamine/Xylazine cocktail. The spinal cord was dissected and then incubated in PFA overnight at 4 °C. Spinal cords were then washed in PBS and incubated in 30% sucrose for cryosectioning. When the spinal cords had sunk in the sucrose solution, they were embedded in Optimal Cutting Temperature (OCT) compound and frozen at −80 °C.

Transverse cryosections (16 or 30 µm) of the cervical spinal cord were obtained using a CM3050S Leica cryostat. For identification and characterization of monosynaptic cholinergic interneurons, 100 μm thick sections were harvested with a Leica VT1000S vibratome. Sections were incubated in PBS containing 1% bovine serum albumin (BSA) and 0.1-0.5% Triton X100 for 2 hours at room temperature. After blocking/permeabilization, tissue was incubated with primary antibodies for overnight to 72 hours. After primary incubation, slides were washed 3 times with PBS, followed by incubation with secondary antibodies for 2 hours to overnight at room temperature. Finally, slides were washed a further 3 times with PBS, mounted sequentially on Superfrost plus gold glass slides (Thermo Scientific) and let dry before applying the Vectashield Vibrance mounting medium (Vector Laboratories) and cover glasses (VWR).

The following primary antibodies were used in this study: goat anti-ChAT (Sigma, RRID: AB_2079751, 1:300), goat anti-VAChT (Millipore, RRID: AB_2630394, 1:1000), rabbit anti-VAChT (Synaptic Systems, RRID: AB_10893979, 1:500), chicken anti-RFP (Rockland, 600901379, 1:500), rabbit anti-DsRed (Takara Bio, RRID: AB_10013483, 1:1000), rabbit anti-c-Fos (Synaptic Systems, RRID: AB_2905595, 1:1000), goat anti-Scip (Santa Cruz Biotechnology, RRID:AB_2268536, 1:5000), rabbit anti-Pitx2 (1:16000)^43^ and rabbit anti-CTB (Novus Biologicals, NB100-63067, 1:500). Fluorophore-coupled secondary antibodies used were: donkey anti-chicken Alexa Fluor 594 (Sigma, SAB4600094), donkey anti-goat Alexa Fluor 405 (Invitrogen, AB_2890272), donkey anti-goat Alexa Fluor 488 (Invitrogen, A11055), donkey anti-goat Alexa Fluor 647 (Jackson ImmunoResearch, AB_2340437), donkey anti-rabbit Alexa Fluor 488 (Abcam, ab150073), and donkey anti-rabbit Alexa Fluor 555 (Invitrogen, A31572).

### Confocal Microscopy and Image Processing

Confocal microscopy images were captured using a Zeiss LSM800 laser scanning confocal microscope, based on an Axio ‘Observer 7’ inverted microscope, equipped with a 20x 0.8 NA apochromatic objective lens. ZEN (blue edition) software was used for image acquisition. Illumination was provided by 405, 488, 555/561, 647 nm laser lines. Unidirectional laser scanning was performed on each channel and images were captured with 8-bit resolution with Gallium Arsenide Phosphid (GaAsP) PMT detectors. Images of PMNs, identified using CTB, were captured from serial Z-stacks with 0.2 µm interval and XYZ voxel dimensions of 312 x 312 x 600 nm, from which resultant 2D images were produced by summating the intensity across 2 µm thick z-stacks passing from the center of the neurons. Images were visualized and processed using FIJI^96^.

### *In situ* hybridization

*In situ* hybridization was performed as previously described^97^. Briefly, a T7 polymerase promoter sequence was added to the 5’ end of the reverse primer of the PCR primers for rabies virus G protein (Forward: AAAGCATTTCCGCCCAACAC, Reverse: TAATACGACTCACTATAGGGCCTCGTCACCGTCCTTGAAA) and DNA was amplified from a plasmid expressing rabies virus G protein (Addgene #26197). Next, RNA probe was generated using T7 polymerase and digoxigenin (DIG) labeling mix. 16 μm-thick cryostat sections from P4 *ChAT::Cre; RphiGT* mice were used for hybridization. Anti-DIG antibody (Roche, RRID: AB_514497) was applied to visualize the signal from the RNA probe.

### Three-dimensional (3D) monosynaptic mapping reconstruction

Images of sections from the brainstem to the spinal cord were organized in sequential order in a folder and imported in Imaris (Oxford Instruments), which automatically generates a 3D brainstem-spinal cord model based on imported sequential images. The contour of the brainstem to spinal cord was outlined by the “Surface” function in Imaris. mCherry+ monosynaptic inputs (magenta) and starter PMNs (turquoise) were identified by the “Spot” function in Imaris and their colors were assigned based on their identities. Starter PMNs are located within clustered ChAT+ MNs in the ventral spinal cord (Extended Data Video 1).

### Filament and VAChT+ synapse analysis

Dendritic morphology of ChAT+ INs was reconstructed and analyzed using the ‘Filament’ function in Imaris. Definitions of the statistics used for filament analysis can be found in the Imaris V 6.3.1 Reference Manual (http://www.bitplane.com/download/manuals/ReferenceManual6_3_1.pdf).

For VAChT+ puncta quantification on the retrogradely traced mCherry+ PMNs, VAChT+ puncta were counted by the ‘Spot’ function in Imaris. Traced mCherry+ PMNs were reconstructed by the ‘Filament’ function in Imaris. Only the VAChT+ puncta that were on the PMNs were included (filtered by the intensity of mCherry+ signal). To categorize subcellular location of the VAChT+ puncta, the cell body of the PMNs was delineated by the ‘Surface’ function in Imaris. VAChT+ puncta that are outside a 100μm radius of the center of the cell body were considered to be on the distal dendrites.

For quantification of puncta colocalization, both VAChT+ and *Pitx2^tdTom^*+ puncta on PMNs were identified by the ‘Spot’ function in Imaris. VAChT+ puncta on the PMNs were then classified as *Pitx2^tdTom^*+ or *Pitx2^tdTom^*-based on the intensity of tdTomato signal in the ‘Spot’ function. The total number of PMNs was counted to calculate the average number of VAChT+ puncta per PMN.

### Topographical analysis

To plot the topographical distribution of mCherry+ ChAT+ INs, the X and Y coordinates of ChAT+ INs, the spinal cord outline, the central canal and MNs were identified in Imaris using the ‘Spot’ function. XY coordinates are rotated and standardized to a spinal cord coordinate map where the central canal is the origin (0, 0). The sizes of the standardized coordinate map are determined by averaging the size of the spinal cord. Since the shape of the spinal cord changes along the rostrocaudal axis and LMNs are located slightly caudal to PMNs, the standardized dimensions of the spinal cord for the ChAT+ INs → PMNs are 2000 μm in the horizontal and 1500 μm in the vertical direction (± 1000 for X axis and ± 750 for Y axis), while the standardized dimensions of the spinal cord for the ChAT+ INs → LMNs are 2000 μm (horizontal) and 1250 μm (vertical) (± 1000 for X axis and ± 625 for Y axis). The absolute values of the X coordinates of these ChAT+ INs were used as their distance to the central canal.

For rostral to caudal distribution of ChAT+ INs that project to PMNs, sections from different animals were aligned based on the spinal cord atlas^98^ and each section was assigned a rostro-caudal position ID. Their XY positions in the spinal cord were determined as described above.

### *In vitro* Isolated Brainstem Spinal Cord Preparation

17 C57/BJ6 neonatal mice (P2-P4) of both sexes were used for *in vitro* electrophysiology experiments. Animals were deeply anesthetized with 4% isoflurane in 100% oxygen before being decerebrated, eviscerated and pinned ventral side down in a dissecting chamber lined with Sylgard silicone elastomer (Dow) that was filled with carbogen-bubbled (95% oxygen, 5% carbon dioxide) artificial cerebrospinal fluid (aCSF, containing 120 mM NaCl, 3 mM KCl, 1.25 mM NaH_2_PO_4_, 1 mM CaCl_2_, 2 mM MgSO_4_, 26 mM NaHCO_3_ and 20 mM D-glucose) at 4 °C. The brainstem and spinal cord were exposed and dissected as previously described^99^, and the brainstem was transected at the pontomedullary junction. Preparations were then transferred to a recording chamber perfused with recirculating, recording aCSF warmed to 25-28 °C and given 1-hour recovery time prior to the initiation of baseline measurements. Extracellular neurograms were obtained using tight fitting suction electrodes attached to the ventral root of the third or fourth cervical spinal segment (C3/4). For split-bath experiments, the recording chamber was divided into two compartments with the use of a plastic wall and by applying Vaseline around the preparation. To confirm successful splitting of the compartments, food coloring was applied sequentially in both compartments at the end of each experiment. Experiments in which any leak of dye was observed from one compartment to the other were excluded from analyses. Signals were amplified 1000 times, and bandpass filtered (10-1000 Hz) using a differential AC amplifier (Model 1700, A-M Systems), digitized at 5 kHz using a Digidata 1440 (Molecular Devices), acquired using Axoscope software (Molecular Devices) and stored on a computer for offline analysis. Signals were analyzed using the Dataview software (courtesy of Dr. W.J. Heitler, University of St Andrews).

### Whole-cell patch clamp electrophysiology

Whole-cell patch clamp recordings were obtained from 22 tdTomato+ interneurons on brainstem-spinal cord preparations obtained from 5 *Pitx2^tdTom^*neonatal mice (P3-P4) of both sexes. Access was gained to Pitx2+ interneurons located near the central canal by performing a mid-sagittal hemisection of the spinal cord using an insect pin. Preparations were stabilized in a recording chamber by pinning them to an agar block and visualized with a 40x objective using infrared illumination and differential interference contrast (DIC) microscopy. Cells were visualized and whole-cell recordings obtained under DIC using pipettes (L: 100 mm, OD: 1.5 mm, ID: 0.84 mm; World Precision Instruments) pulled on a Flaming Brown micropipette puller (Sutter Instruments Model P97) to a resistance of 2.5-3.5 MΩ. Pipettes were back-filled with intracellular solution (containing 140 mM KMeSO4, 10 mM NaCl, 1 mM CaCl2, 10 mM HEPES, 1 mM EGTA, 3 mM Mg-ATP and 0.4 mM GTP-Na2; pH 7.2-7.3, adjusted with KOH). Signals were amplified and filtered (6 kHz low pass Bessel filter) with a Multiclamp 700B amplifier, acquired at 20 kHz using a Digidata 1440A digitizer with pClamp Version 10.7 software (Molecular Devices) and stored on a computer for offline analysis.

All interneuron intrinsic properties were studied by applying a bias current to maintain the membrane potential at −60 mV. Values reported are not liquid junction potential corrected to facilitate comparisons with previously published data^43,45^. Cells were excluded from analysis if access resistance was greater than 20 MΩ, changed by more than 5 MΩ over the duration of the recording, or if spike amplitude was less than 60 mV when measured from threshold (described below).

Basal passive properties including capacitance (C), membrane time constant (tau), and input resistance (Ri) were measured during a hyperpolarizing current pulse that brought the membrane potential from −60 to −70mV. Input resistance was measured from the initial voltage trough to minimize the impact of active conductances with slow kinetics (eg. Ih, sag). The time constant was measured as the time it took to reach 2/3 of the peak voltage change. Capacitance was calculated by dividing the time constant by the input resistance (C = tau/Rin). Resting membrane potential was measured, from the MultiClamp Commander, 10 minutes after obtaining a whole-cell configuration.

Rheobase was measured using short (10 ms) depolarizing current steps and determined as the first current step in a series of three steps to elicit an action potential. The voltage threshold of the first action potential was defined as the voltage at which the change in voltage reached 10 mV/ms. The amplitude of synaptic currents and the frequency of current amplitude and action potentials were measured during respiratory bursting using a threshold-based event detection in Clampfit (Molecular Devices).

### Adult Perfused Preparation

9 prepubescent male Sprague Dawley rats (age: 4-6 weeks; 80-180g) were deeply anesthetized with 5% isoflurane in air before being bathed in ice-chilled physiologic saline solution (115 mM NaCl, 4 mM KCl, 1 mM NaHCO3, 1.25 mM NaH2PO4, 2 mM CaCl2, 10 mM D-glucose, and 12 mM sucrose), decerebrated at approximately the mid collicular level, and spinally transected near the thoracolumbar junction. Rats were eviscerated and vagotomised, and then perfused via the descending aorta with physiologic saline solution equilibrated to 40 mmHg PCO2 and balance oxygen, pressure held above 90 mmHg, and the temperature at 32-33 °C. Extracellular neurograms were obtained from the phrenic nerve using silver wire hook electrodes and electromyograms recorded from muscles of the 5th intercostal muscles. Signals were acquired at 5 kHz, amplified 1000 times, and bandpass filtered from 0.1 to 1 kHz (Axoscope 9.0). Pharmacological manipulations of M2 receptors were performed by delivering 10 µM methoctramine (Sigma-Aldrich, M105) through the perfusate.

### Whole body plethysmography

Freely-moving mice were placed in a chamber for whole body plethysmography (emka, Fig. 7A). The air flow was maintained at 0.75 L/min per chamber for all gas mixtures. Breathing measurements were obtained from pairs of adult mice (P60-120), with each pair consisting of one control and its sex-matched mutant littermate. Normal air was given for 45 minutes, followed by 5% or 10% CO_2_ for 15 minutes (Fig. 7A). All breaths were collected initially and plotted for overview. Since breathing patterns can greatly vary under normal air depending on the activity level of the animal, we selected breaths during which the animal was resting (not moving, sniffing or grooming, accompanied by a consistent pattern of low frequency breaths) to represent breathing under normal air conditions. When switching from normal air to hypercapnic conditions, a response curve was observed due to the gas exchange in the chamber being a gradual process. We selected breaths after maximum breathing frequency was reached after the switch to represent breaths under hypercapnic conditions. For each animal, at least 3 trials were performed, and all the trials were included in the analysis. For *Dbx1^ΔChAT^* mice and their control littermates, 8697-30757 breaths under air and 11793-68143 breaths under 10% CO_2_ for each animal were included for analysis. For *Pitx2^ΔChAT^* mice and their control littermates, 997-28228 breaths under air, 6727-48128 breaths under 10% CO_2_ and 2787-22118 breaths under 5% CO_2_ for each animal were included for analysis. All qualifying breaths were used to characterize their distribution. The mean values from all qualifying breaths collected were used to represent individual animals for group comparisons. Normalization was presented as fold control, where the control was the matched littermate that was recorded at the same time.

### Hypercapnic challenge and c-Fos expression analysis

P20 *ChAT::eGFP* mice were placed in whole body plethysmography chambers for normal air or hypercapnic gas challenge. Either normal air (79% N_2_, 21% O_2_), 5% CO_2_ (with 74% N_2_ and 21% O_2_), or 10% CO_2_ (with 69% N_2_ and 21% O_2_) were given for either 1 hour or for 15 minutes. The 15-minute trials were followed by 45 minutes of normal air. Mice were euthanized by intraperitoneal injection of a ketamine/xylazine cocktail solution and dissected immediately after the gas challenge. Continuous spinal cord sections were mounted on 10 individual slides (10 replicates). One set of the spinal cord sections was used for c-Fos immunostaining and quantification. The numbers shown in Fig. 6 and Extended Data Fig. 6 are total numbers from one set of spinal cord sections and correspond to one tenth of the overall PMN/ChAT+ IN numbers in one animal. c-Fos+ and ChAT+ cells were identified separately by using the ‘Spot’ function in Imaris. c-Fos+ cells in the white matter of the spinal cord were rare and were not included in our study. Based on their location in the spinal cord, neurons were divided into MN and non-MN groups. To define c-Fos/ChAT colocalization and count c-Fos+ ChAT+ INs, a filter named “shortest distance to ChAT+ non-MN spots” was applied to c-Fos+ non-MN spots.

### Experimental design and statistical analysis

For rigor and reproducibility, both male and female mice were used for all reported results. Data were presented as box plots or bar plots with each dot representing data from an individual mouse. Violin plots were used for the morphological comparison of ChAT+ interneurons for better demonstration of the distribution. For violin plots, solid lines indicate the mean and dashed lines indicate 25^th^ and 75^th^ percentile. Two-factor repeated measures ANOVA were conducted to test the effect of pharmacological agents on intrinsic properties and currents, with MN subtype and drug as factors. Appropriate and equivalent nonparametric tests (Mann-Whitney or Kruskal-Wallis) were conducted when data failed tests of normality or equal variance with Shapiro Wilk and Brown-Forsythe tests, respectively. Paired or unpaired t-tests were performed on data with two variables. One sample t-test (hypothetical value = 1) was used for data after normalization. Individual data points for all recorded *Pitx2^Tdtom^*+ cells are presented in figures along with mean ± SD, and were reported in Extended Data Table 1. Statistical analyses were performed using Graph Pad Version 9.0 (Prism, San Diego, CA, USA). p < 0.05 was considered to be statistically significant, where *p < 0.05, **p < 0.01, ***p < 0.001, and ****p < 0.0001.

## Acknowledgements

We thank Steven Crone, Britton Sauerbrei and Rishi Dhingra for helpful discussions and comments on the manuscript, Niccolo Zampieri for providing *Pitx2::Cre* mice and Susan Brenner-Morton for the Pitx2 antibody. This work was funded by NIH R01NS114510 to PP, F31NS124240 to MTM, a Tenovus Scotland grant to GBM and SAS and CIHR RN387354 to RJAW. PP is the Weidenthal Family Designated Professor in Career Development.

