## Extended Data for "A cholinergic spinal pathway for the adaptive control of breathing"

### Extended Data Figures

**Extended Data Figure 1. Validation of monosynaptic retrograde rabies virus tracing strategy**

(A) Genetic schematic of *ChAT::Cre; RphiGT* mice. *TVA protein was reported to be predicted as non-functional due to a 1bp insertion immediately following the TVA start codon.

(B) Expression of rabies *G protein* (*in situ* hybridization) was only detected in MNs (ChAT, green) in *ChAT::Cre; RphiGT* mice at P4. Scale bar = 100 µm.

(C) Specific unilateral labeling of PMNs after diaphragm injections. Only ventral roots at cervical (from PMNs), but not thoracic or lumbar, spinal cord levels were labeled.

**Extended Data Figure 2. Monosynaptic retrograde tracing of PMNs**

(A) mCherry-labeled PMN after unilateral injection of RabiesΔG-mCherry into the diaphragm (starter PMN). Scale bar = 200 µm and 50 µm (inset).

(B) Quantitation of starter PMNs in individual animals (n = 7).

(C) Brainstem PMN inputs are evenly distributed between the contralateral and ipsilateral injection sides.

(D and E) Connectivity index for total monosynaptic input, brainstem input, spinal cord input, and ChAT+ IN input in individual animals (n = 7).

(F and G) Examples of PMN-projecting ChAT- interneurons from the ventral (F) and dorsal (G) spinal cord at cervical and thoracic levels. Scale bar = 200 µm (top) and 50 µm (bottom).

(H) In the intermediate spinal cord, ~50% of inputs (mCherry+ cells excluding starter PMNs) are ChAT+ INs.

(I) Percentage of inputs derived from ChAT+ INs in the entire spinal cord and at cervical, brachial and thoracic levels.

**Extended Data Figure 3. Morphological comparison of contralateral and ipsilateral ChAT+ INs → PMNs**

(A)  Morphological analysis of a single traced ChAT+ IN → PMNs.

(B-C) Sholl analysis of contralateral and ipsilateral ChAT+ INs → PMNs. Contralateral ChAT+ INs → PMNs have more maximum Sholl intersections than ipsilateral ChAT+ INs → PMNs (C).

(D-G) Contralateral and ipsilateral ChAT+ INs → PMNs have similar overall dendritic length (D), area (E), maximum branch level (F) and maximum branch depth (G).

* $p < 0.05$

**Extended Data Figure 4. Distribution of cholinergic synapses on a single retrogradely traced PMN**

(A-B) Cholinergic synapses (VAChT+ puncta, green) on a single retrogradely traced PMN (RabiesΔG-mCherry, red) in a control mouse (P11). Scale bar = 100 µm.

(C-E) Enlargement of square region in (A). VAChT+ puncta on the traced PMN were labeled by spots (Imaris software). Dendrites within 100 µm radius are considered as proximal while the rest are considered as distal. White arrows indicate VAChT+ puncta on distal dendrites (purple). Square region in (C) was enlarged in (D-E). VAChT+ puncta on the MN soma and proximal dendrites were color-coded in yellow and turquoise, respectively. Scale bar = 20 µm (C) and 5 µm (E).

(F) Distribution of VAChT+ puncta on retrogradely traced PMNs (n = 6 cells).

**Extended Data Figure 5. Validation of *Pitx2*$^{tdTom}$ mice**

(A) Representative image of Pitx2 immunostaining (green) in the cervical spinal cord of e16.5 *Pitx2*$^{tdTom}$ mice. PMNs are labeled with Scip (magenta).

(B) Over 90% of Pitx2+ cells were tdTomato+ in *Pitx2*$^{tdTom}$ mice.

(C) Average number of cholinergic synapses (VAChT+) on PMN somas during early postnatal development (n = 3 per group).

(D) Percentage of VAChT+ puncta that are *Pitx2*$^{tdTom}$+ on PMN somas during early postnatal development (n = 3 per group).

**Extended Data Figure 6. c-Fos activation in the spinal cord under a hypercapnic gas challenge**

(A) Number of spinal cord c-Fos+ cells (excluding MNs) after 10% $CO_2$ for 1 hour.

(B-C) Mean intensity levels of c-Fos+ PMNs were significantly elevated after 10% $CO_2$ for 1 hour (B). Distribution of mean intensity from all c-Fos+ MNs in individual animals (C).

(D-E) Mean intensity levels of c-Fos+ ChAT+ INs were comparable to baseline after 10% $CO_2$ for 1 hour (D). Distribution of mean intensity from all c-Fos+ ChAT+ INs in individual animals (E).

**Extended Data Figure 7. Validation of mouse models for blocking cholinergic neurotransmission from $V0_C$ interneurons**

(A) Genetic schematic of *Dbx1*$^{\Delta ChAT}$ and *Pitx2*$^{\Delta ChAT}$ mice.

(B-C) Expression of ChAT (green) in VAChT+ puncta (red) on PMNs in *Dbx1*$^{\Delta ChAT}$ and control mice. Numbered synapses in (B) were enlarged in (C). Scale bar = 20 µm (B, left), 5 µm (B, right), and 1 µm (C).

(D) Percentage of ChAT+ VAChT+ puncta on PMNs in *Dbx1*$^{\Delta ChAT}$ and control mice. 10 cells from each animal were quantified (n = 3 per group).

(E) Number of VAChT+ puncta on single PMNs in *Dbx1*$^{\Delta ChAT}$ and control mice. 10 cells from each animal were quantified (n = 3 per group).

(F-G) Expression of ChAT (green) in VAChT+ puncta (red) on PMNs in *Pitx2*$^{\Delta ChAT}$ and control mice. Numbered synapses in (F) were enlarged in (G). Scale bar = 20 µm (F, left), 5 µm (F, right), and 1 µm (G).

(H) Percentage of ChAT+ VAChT+ puncta on PMNs in *Pitx2*$^{\Delta ChAT}$ and control mice. 10 cells from each animal were quantified (n = 3 per group).

(I) Number of VAChT+ puncta on single PMNs in *Pitx2*$^{\Delta ChAT}$ and control mice. 10 cells from each animal were quantified (n = 3 per group).

**Extended Data Figure 8. Cholinergic interneuron silencing impairs the response to hypercapnia**

(A-B) Examples of breath traces under normal air and 10% $CO_2$ in *Dbx1*$^{\Delta ChAT}$ and their control littermates. A single breath was enlarged in (B).

(C, E, G, I, and K) Breath frequency (C), minute ventilation (E), tidal volume (G), PIF (I), and PEF (K) distribution, and their normalized value under normal air in *Dbx1*$^{\Delta ChAT}$ and control mice (n = 5 per group).

(D, F, H, J, and L) Frequency (D), minute ventilation (F), tidal volume (H), PIF (J), and PEF (L) distribution and their normalized value under 10% $CO_2$ in *Dbx1*$^{\Delta ChAT}$ and control mice (n = 5 per group.

(M-N) PEF distribution and normalized value under normal air (M) and 10% $CO_2$ (N) in *Pitx2*$^{\Delta ChAT}$ and control mice (n = 14-17 per group).

**Extended Data Figure 9. Cholinergic interneuron silencing does not alter the response to moderate hypercapnia (5% $CO_2$)**

(A-B) Examples of breath traces under normal air and 5% $CO_2$ in *Pitx2*$^{\Delta ChAT}$ and control mice. A single breath was enlarged in (B).

(C, E, G, I, and K) Frequency (C), minute ventilation (E), tidal volume (G), PIF (I), and PEF (K) distribution under 5% $CO_2$ in *Pitx2*$^{\Delta ChAT}$ and control mice (n = 14-17 per group).

(D, F, H, J, and L) Mean and normalized frequency (D), minute ventilation (F), tidal volume (H), PIF (J), and PEF (L) under 5% CO2 condition in *Pitx2*$^{\Delta ChAT}$ and control mice.

**Extended Data Video 1. Three-dimensional (3D) monosynaptic mapping reconstruction**

3D reconstruction of mCherry+ PMN monosynaptic inputs (magenta) in the brainstem and spinal cord in a single animal. Starter PMNs are indicated in turquoise.

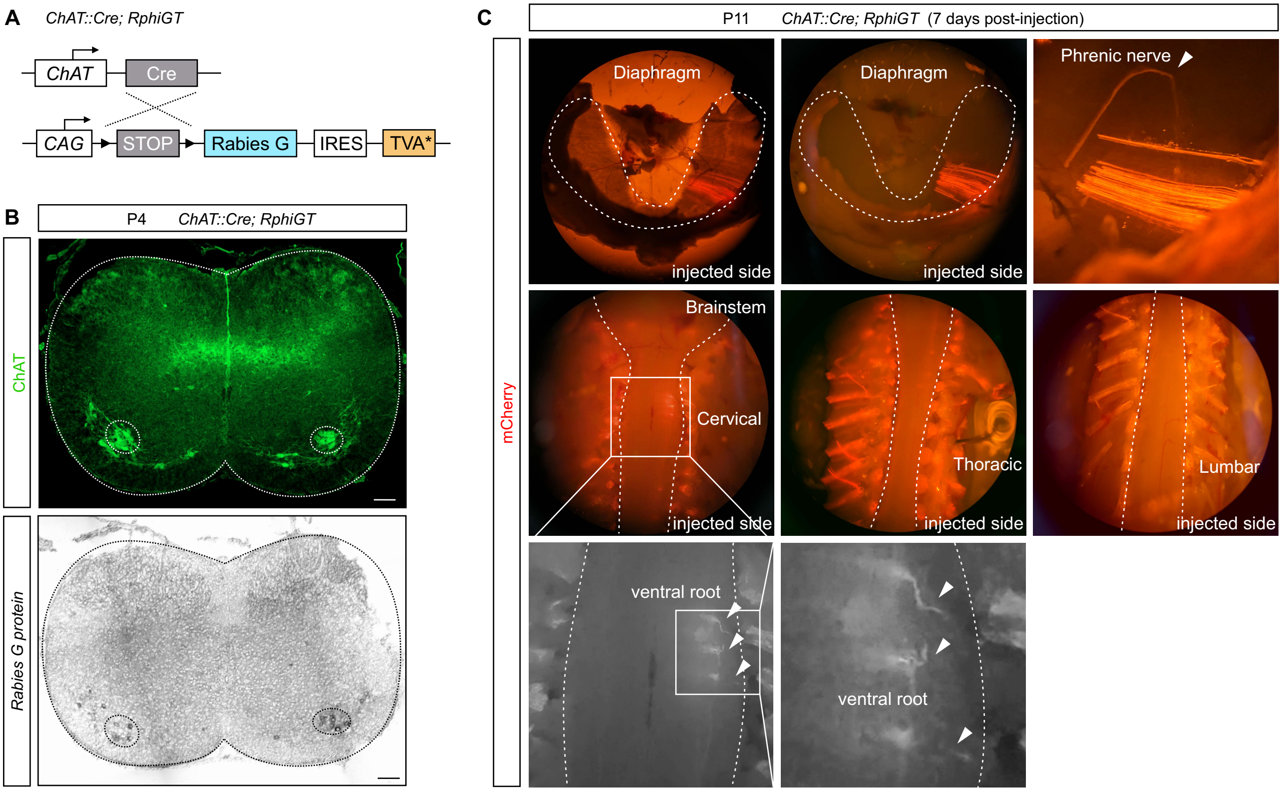


Extended Data Figure 1

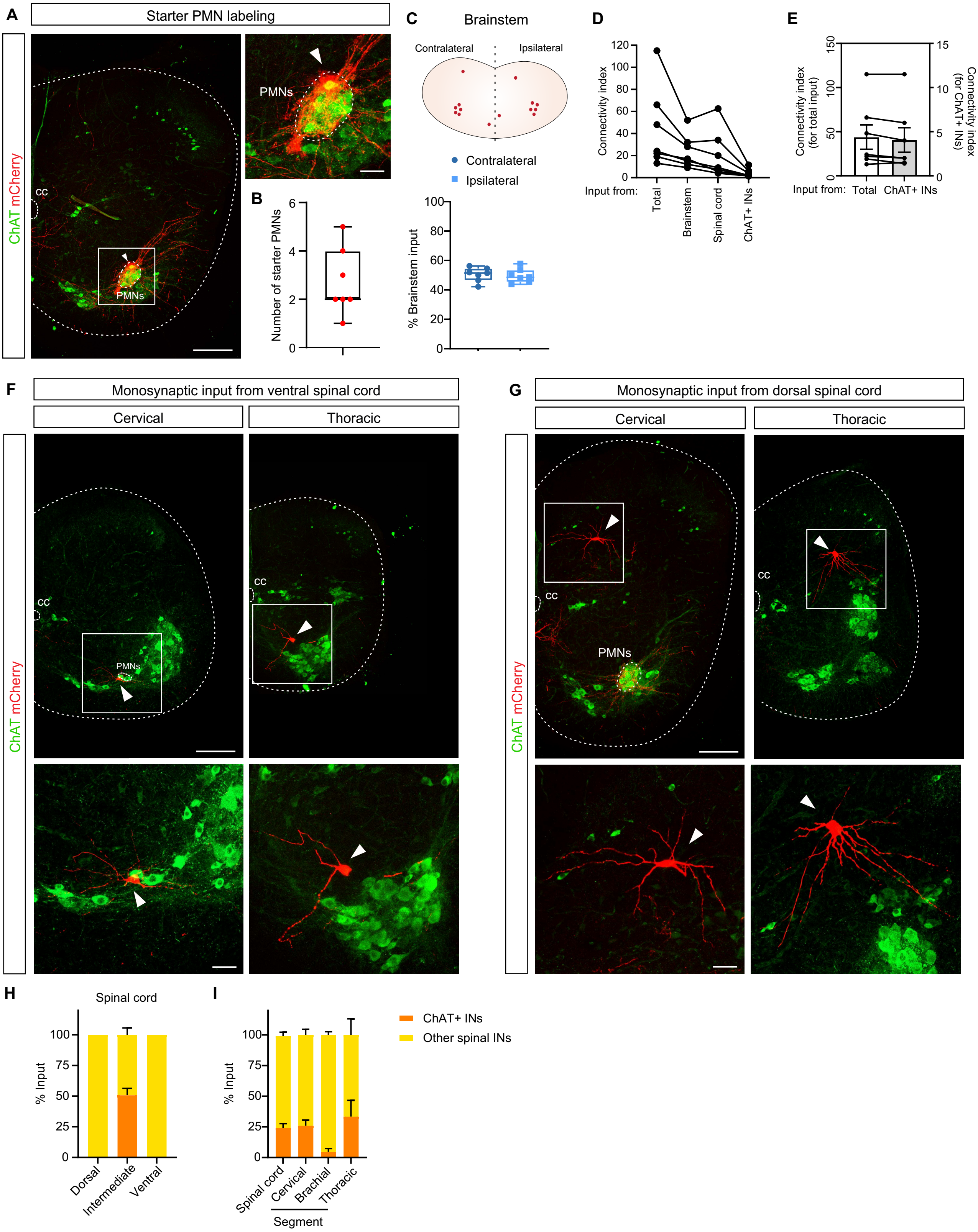


Extended Data Figure 2

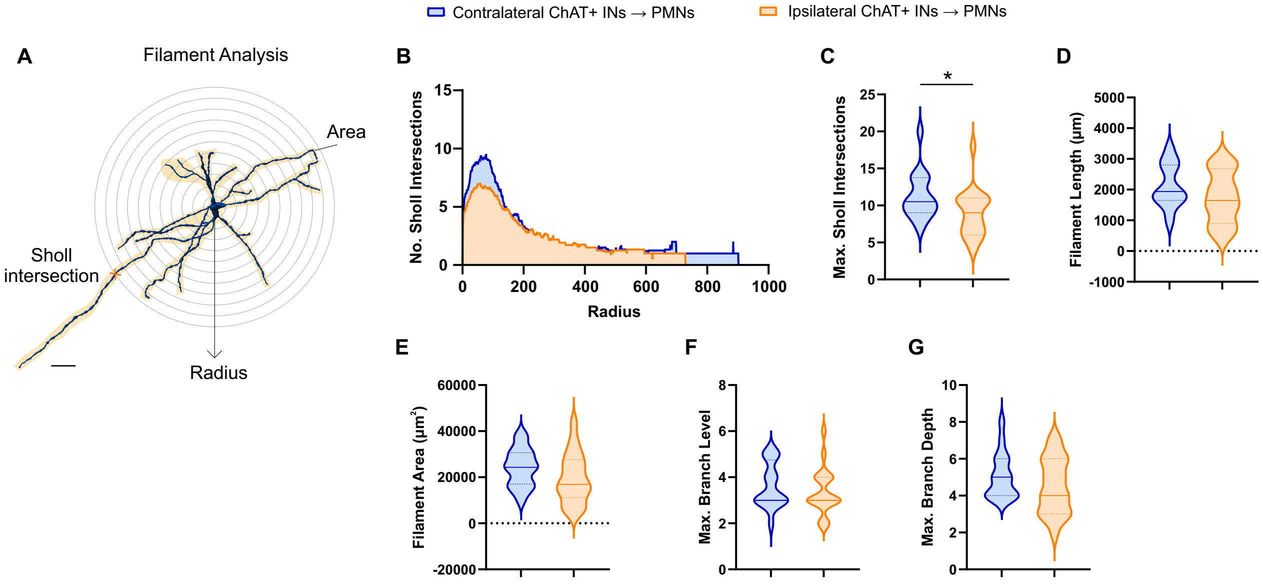


Extended Data Figure 3

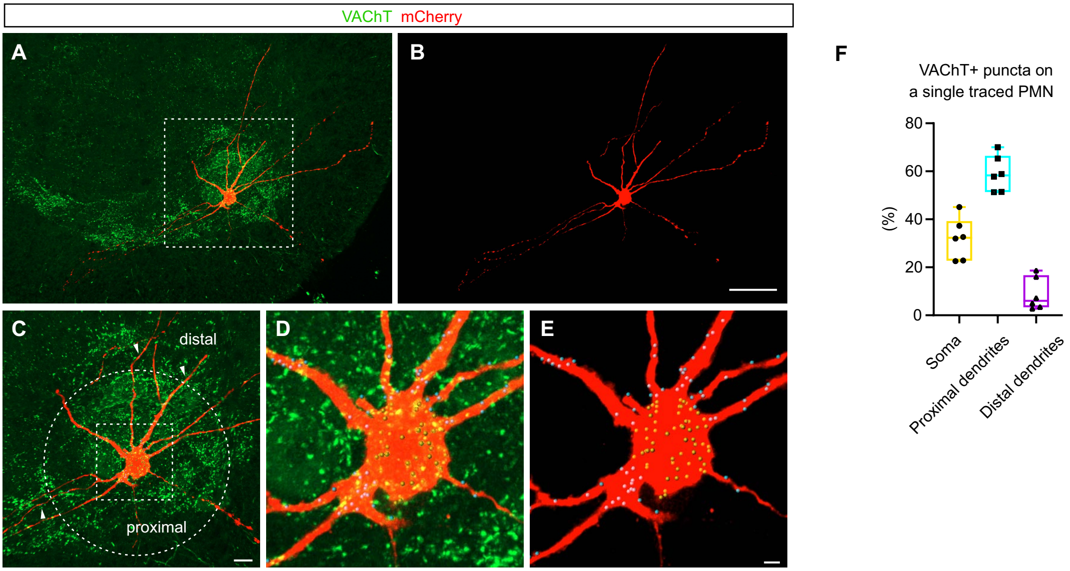


Extended Data Figure 4

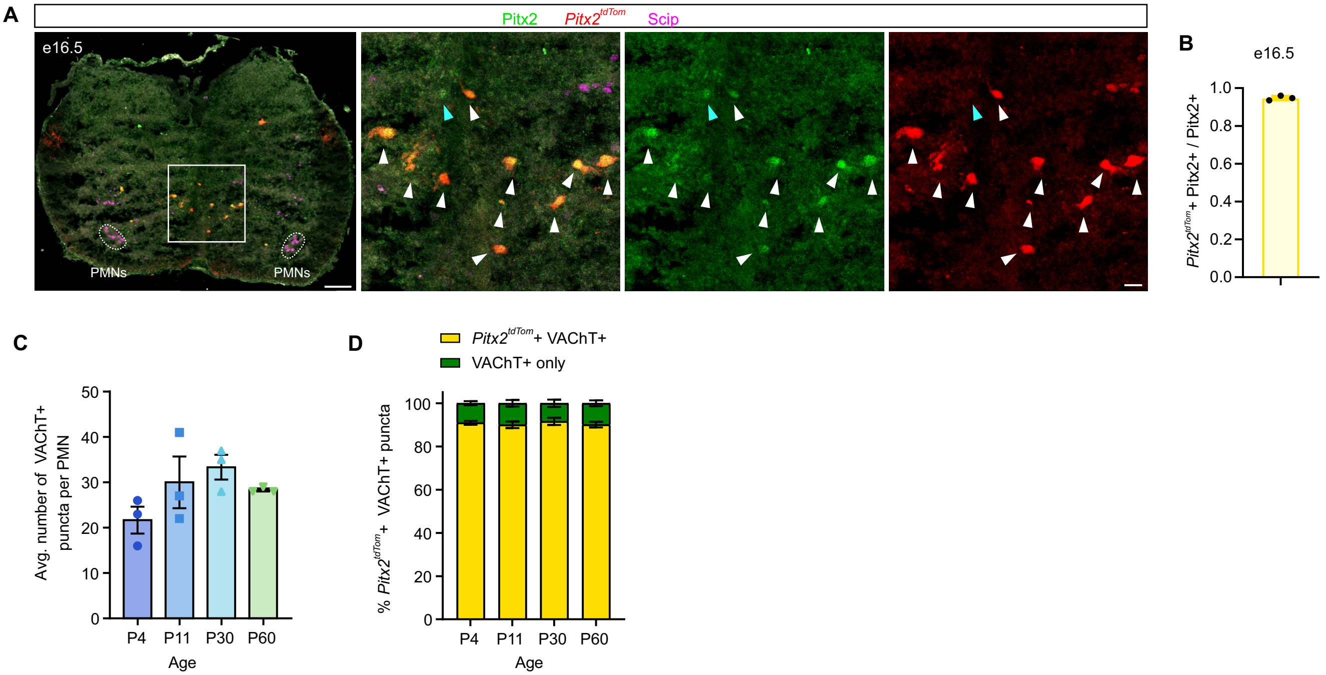


Extended Data Figure 5

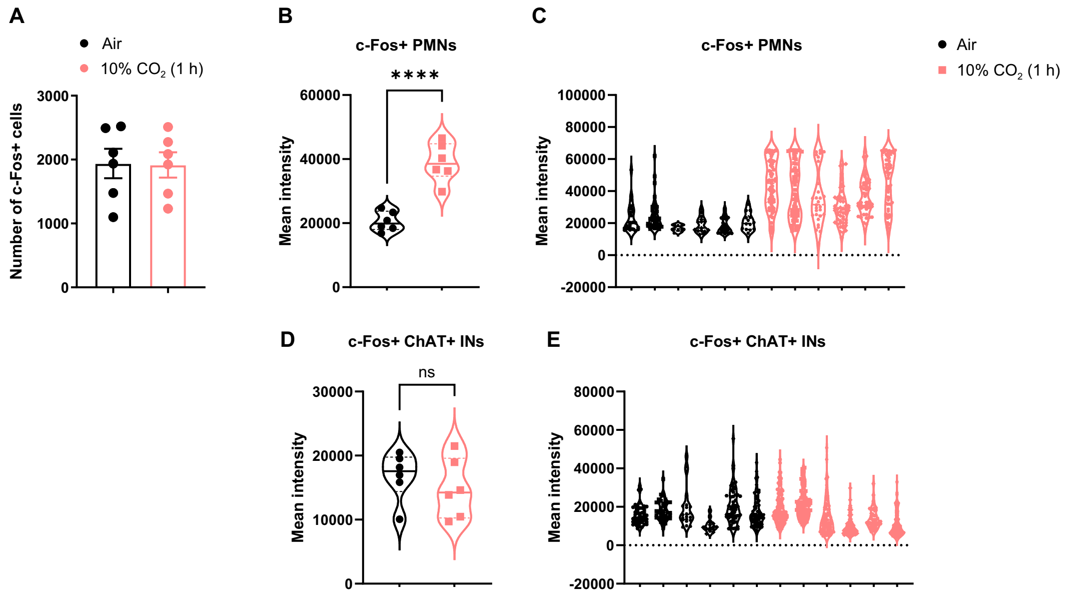


Extended Data Figure 6

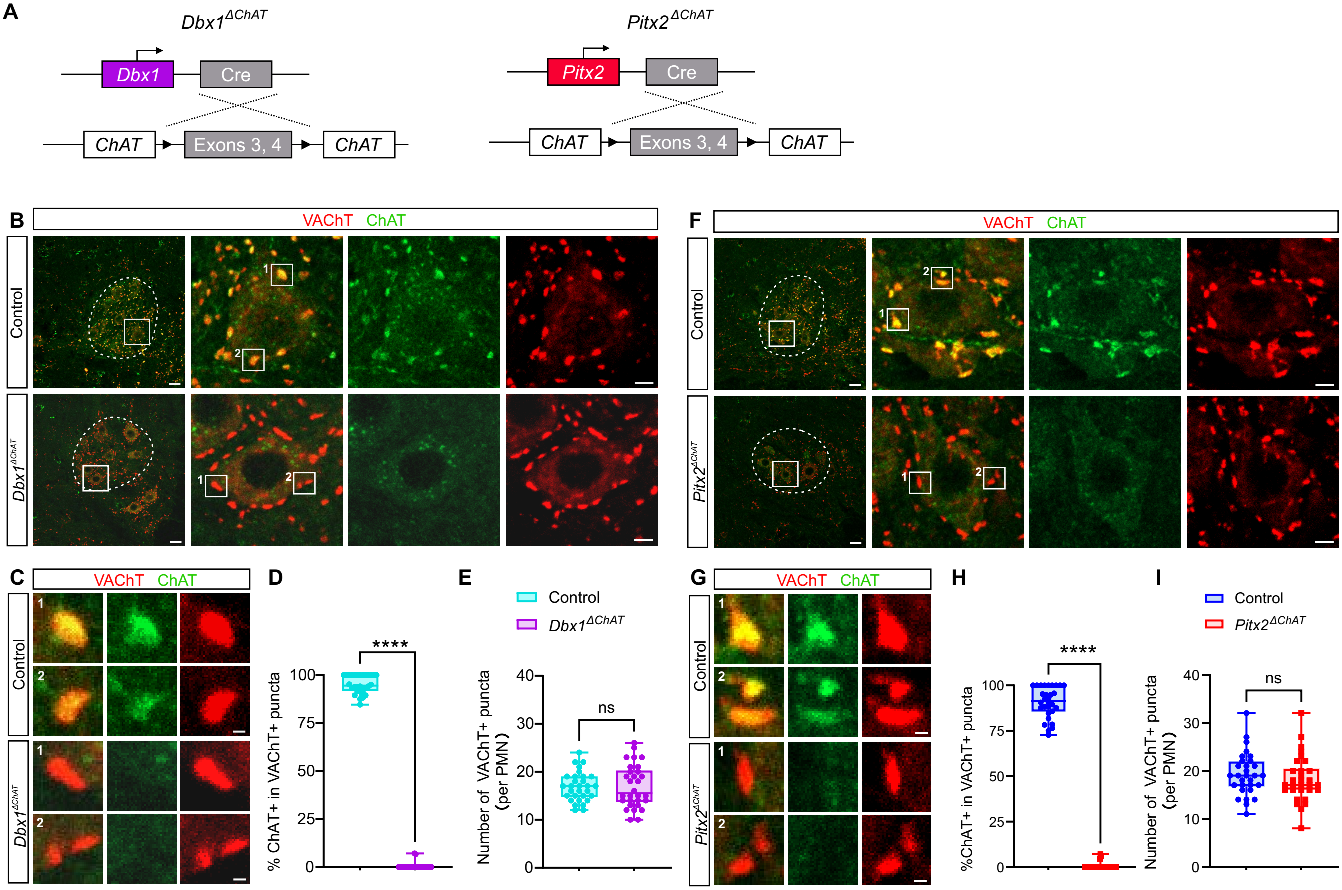


Extended Data Figure 7

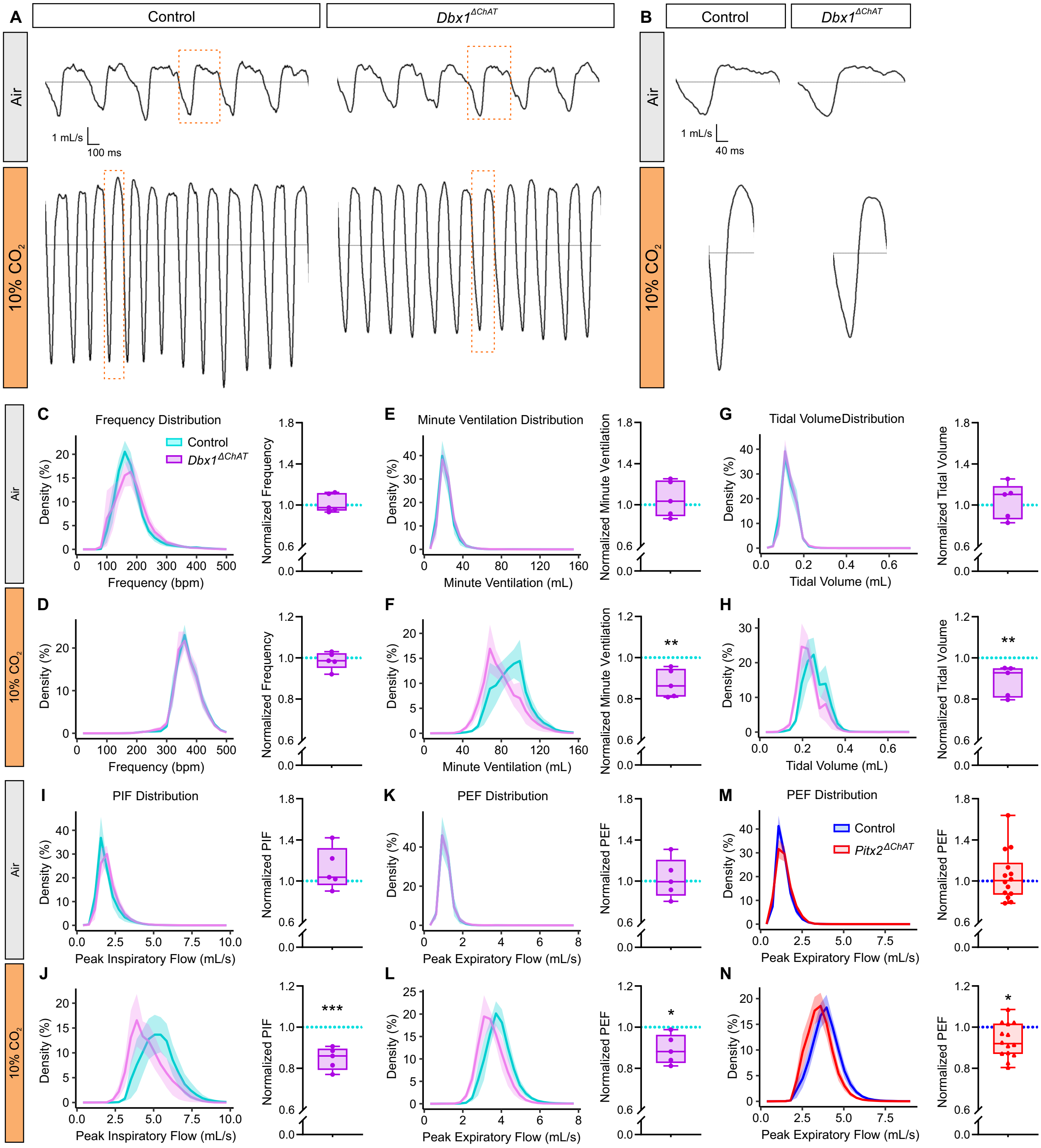


Extended Data Figure 8

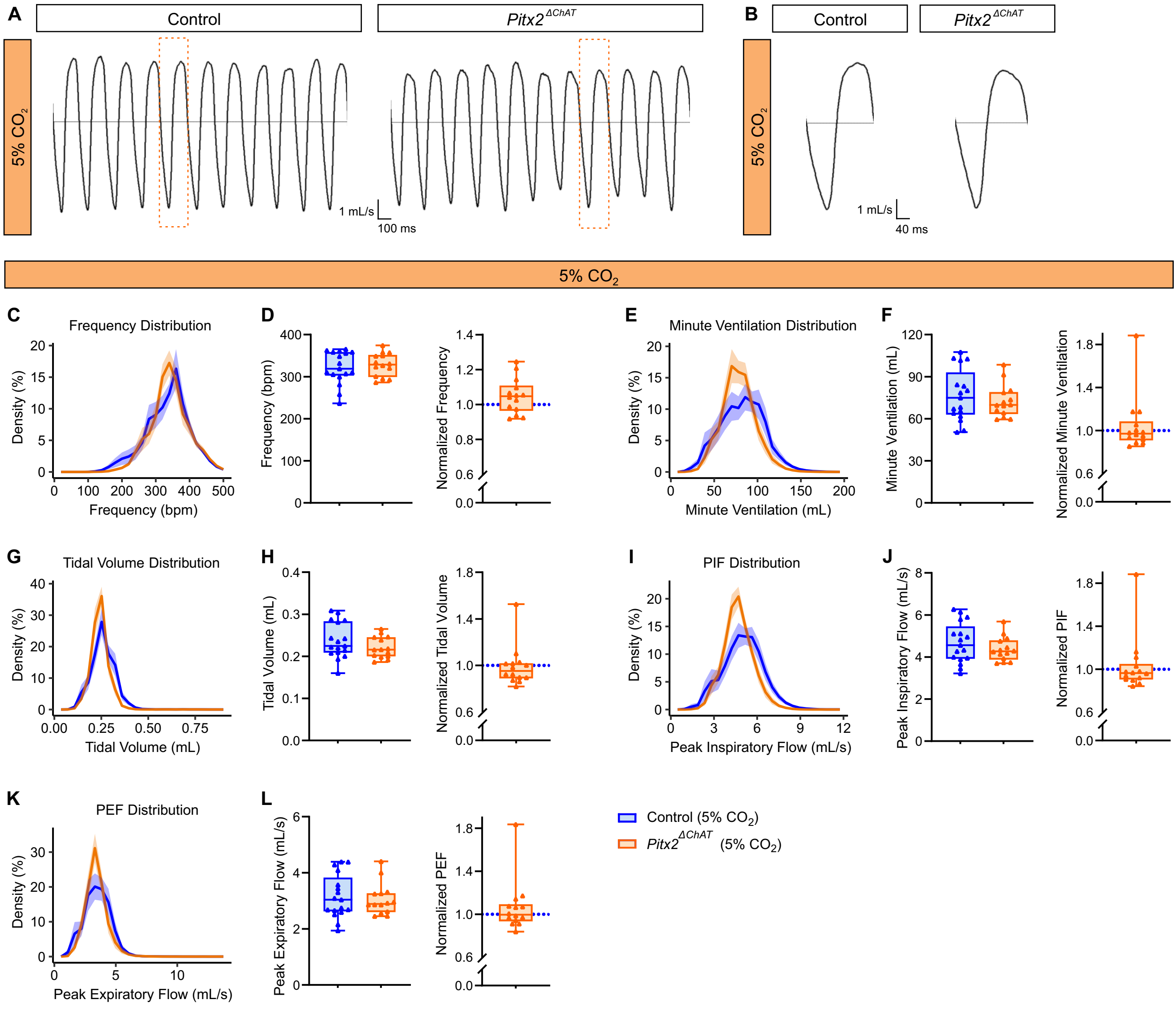


Extended Data Figure 9

Extended Data Table 1: Intrinsic properties of respiratory-related and non-respiratory related *Pitx2*$^{tdTom}$+ interneurons in the C3/C4 spinal segments.

| | Respiratory Related (n = 4) | Non-Respiratory Related (n = 4) | U | p |
|---|---|---|---|---|
| Capacitance (pF) | 132±63 | 107±6.7 | 6 | 0.7 |
| Tau (ms) | 78.3±43.5 | 79.1±4.3 | 8 | >0.99 |
| Input Resistance (MOhm) | 587±81 | 822±313 | 4 | 0.3 |
| Resting Membrane Potential (mV) | -58.6±4.9 | -56.1±6.7 | 6 | 0.7 |
| Rheobase (pA) | 136±19 | 142±61 | 8 | >0.99 |
